# An updated roadmap for brain-mental health associations in the UK Biobank

**DOI:** 10.64898/2026.09.09.750514

**Authors:** Xuqian Li, Sarah Khalife, Lena K L Oestreich

**Author notes:** **Correspondence:** Xuqian Li, Australian Institute for Bioengineering and Nanotechnology, Building 75, Cnr College Rd &, Cooper Rd, St Lucia QLD 4067, Australia.

## Abstract

The UK Biobank offers an unprecedented opportunity to examine associations between brain imaging and mental health at scale, but their magnitude and replicability remain unclear. We correlated 17,580 imaging-derived phenotypes across seven MRI modalities with 309 mental health phenotypes in 8,415 participants, split into discovery and validation subsets. Associations were generally small (median |*r*| = 0.018; top 1% of 0.108), and they replicated poorly: only 9.0% of significant discovery associations were significant in validation. Eating disorders showed the largest effects of any mental disorders but replicated in only 6.0% of cases, whereas substance use showed the highest replication (14.5%). Susceptibility-weighted imaging produced the most replicable imaging associations. Longitudinal changes in brain and mental health measures were weakly related. We provide an interactive dashboard that returns the effect-size and replicability benchmarks for any combination of mental health category and imaging modality, supporting variable selection and study design in UK Biobank research.

## 1 Introduction

Mental disorders are among the leading contributors to disability worldwide and affect a substantial proportion of the population across the lifespan (GBD 2019 Mental Disorders Collaborators, 2022). Identifying the neurobiological correlates of these conditions has long been a central aim of psychiatric neuroscience, on the expectation that measurable features of brain structure and function will help explain who develops mental illness and why. For most of its history, this work has relied on clinical neuroimaging studies with small samples and narrow inclusion criteria. Such studies have produced brain-mental health associations that are often inconsistent from one report to the next and difficult to generalise beyond the specific groups examined (Dutt et al., 2022). Large-scale imaging studies have since offered an explanation for this inconsistency. In a landmark brain-wide association study (BWAS), with a sample size of around 50,000, Marek et al. (2022) showed that univariate correlations between imaging-derived phenotypes (IDPs) and behavioural measures are substantially smaller than the earlier literature had assumed, with typical correlations around 0.01 against the effects of 0.2 or above routinely reported in smaller studies. Effects this small, estimated at the sample sizes common in neuroimaging, are exactly what produce the inflated and poorly replicating findings behind the inconsistency noted above. Interpreting any single finding correctly, and planning a study with the power to detect a real effect, therefore depends on knowing the realistic size and replicability of these associations.

The UK Biobank (UKB) is well suited to provide such benchmarks. It is a prospective cohort of approximately half a million UK adults, aged 40 to 69 at recruitment between 2006 and 2010, followed since through repeat visits, linked health records, and successive questionnaires (Miller et al., 2016). An imaging enhancement launched in 2014 set out to acquire multimodal brain magnetic resonance imaging (MRI) from 100,000 participants, with repeat scans in 60,000, across seven modalities capturing cortical and subcortical structure, white matter microstructure, cerebral perfusion, and spontaneous and task-related brain activity (Littlejohns et al., 2020; Miller et al., 2016). A standardised automated pipeline further processes the raw images into thousands of IDPs, released ready for analysis by the UKB. The UKB also holds a wide range of mental health phenotypes (MHPs), collected through a baseline touchscreen questionnaire and two later online questionnaires that capture current and lifetime symptoms of common disorders, transdiagnostic features, and environmental risk factors (Davis et al., 2020; Davis, Coleman, et al., 2025). Together, this combination of repeated multimodal imaging and transdiagnostic mental health phenotyping supports estimation of brain-mental health associations at a scale sufficient for reliable benchmarking.

Dutt et al. (2022) provided the first such benchmarks in the UKB, estimating effect sizes for five summary scores of depression and anxiety against a dimensionally reduced set of IDPs. That work gave the field its first population-scale reference values for brain-mental health associations in the cohort. Both the mental health assessment and the imaging protocol have expanded substantially since then. On the imaging side, arterial spin labelling (ASL) has been added to the protocol, with its first set of IDPs released in 2023. ASL measures cerebral blood flow, a physiologically distinct signal that structural and functional MRI do not capture and is increasingly used to study neurological and psychiatric disorders (Haller et al., 2016). On the mental health side, the most recent wave of online questionnaires introduced several new topics, including eating disorders (ED), which carry one of the highest mortality rates in psychiatry (Arcelus et al., 2011) and whose structural brain alterations are among the largest seen in any psychiatric condition (Walton et al., 2022). The UKB has also gained a longitudinal dimension following the release of data from the second imaging visit, which began in 2019 and repeats the scan roughly five years after the first. This makes it possible to ask not only how brain and mental health are associated across individuals, but whether they change together within an individual over time. Most of this enlarged space has not been characterised, and how brain and mental health relate across the majority of these measures remains unknown.

The present study provides an updated roadmap and characterisation of brain-mental health associations in the UKB, drawing on data available as of the April 2025 release. We benchmark univariate correlations between 17,580 IDPs from seven imaging modalities and 309 MHPs spanning 15 categories, estimating both the magnitude and replicability of these associations across participants subsets. We further examine how association magnitude and replicability vary across mental health categories, imaging modalities, and combinations of mental health and imaging phenotypes. Using measurements available across two imaging visits, we additionally quantify within-subject longitudinal associations to assess the extent to which changes in brain and mental health measures covary over time. Together, these analyses provide empirical benchmarks for interpreting brain-mental health associations and informing study design and power calculations in the UKB. To facilitate their use, we provide an interactive web application through which researchers can query effect-size and replicability benchmarks across mental health categories and imaging modalities.

## 2 Methods

### 2.1 Dataset

Data were obtained from the UKB (April 2025 release, application number 100773), a prospective epidemiological resource comprising approximately 501,300 participants recruited between 2006 and 2010 at assessment centres across the UK. Participants have since been followed through repeat assessment visits, linked health records, and successive online questionnaires, with an imaging study launched in 2014 (Littlejohns et al., 2020; Miller et al., 2016).

### 2.2 Mental health phenotypes

#### 2.2.1 Data description

Mental health data within the UKB are derived from multiple sources: (a) self-reported responses to touchscreen questionnaires administered at UKB Assessment Centres (<u>Category</u> <u>100025</u>) and to online follow-up questionnaires completed independently of Assessment Centre visits (<u>Category 100089</u>); (b) linkage to various health records produced by the National Health Service (<u>Category 100091</u>); and (c) genetic profiles (<u>Category 100314</u>), including polygenic risk scores for mental disorders (Davis, Mirza, et al., 2025).

The current project focuses specifically on self-report questionnaire data. Overall, the self-report questionnaires in the UKB capture data across three key domains in mental health research, including current and lifetime symptoms and/or diagnosis of common mental disorders, transdiagnostic features, and environmental factors known to be associated with mental disorders (Davis, Mirza, et al., 2025). In the following sections, we provide a brief description of each wave of questionnaire assessment (see **Table S1** for a list of official UKB resources).

As part of the baseline assessment (Initial assessment visit [2006–10]), participants provided informed consent and completed a self-report Touchscreen Questionnaire (T/Screen 2009+), which included several questions related to mental health. These covered areas including current mood, lifetime mental disorder, neuroticism, and social support. In April 2009, the questionnaire was expanded to include additional items that enabled the provisional classification of major depression and bipolar disorder (BD; Smith et al., 2013), along with new questions assessing subjective well-being. Between August 2012 and June 2013, over 103,000 participants were re-invited, and more than 20,000 returned to UKB Assessment Centres to repeat the baseline assessment, including completion of the T/Screen 2009+ for a second time (First repeat assessment visit [2012–13]). In 2014, the UKB imaging study was launched with the goal of re-inviting 100,000 participants to undergo multimodal imaging at UKB Assessment Centres (Imaging visit [2014+]; Littlejohns et al., 2020). During this visit, participants once again completed the T/Screen 2009+. As of April 2025, more than 89,000 participants had completed this imaging visit. In May 2019, the UKB initiated a repeat imaging project, aiming to re-invite 60,000 participants who had completed the initial imaging visit to return to the same Assessment Centres for a second imaging session (First repeat imaging visit [2019+]). The T/Screen 2009+ was again administered during this visit. By April 2025, over 12,000 participants had completed the repeat imaging visit.

In July 2016, to further enrich UKB’s phenotyping of mental disorders, the online Mental Health Questionnaire (MHQ 2016+) was introduced. This questionnaire was designed to capture additional information on current and lifetime symptoms of common mental health conditions, including mood disorders, generalised anxiety, and substance use. It also included new items addressing psychotic experiences, adverse life events, self-harm and suicidal thoughts, and subjective well-being (Davis & Hotopf, 2019; Davis et al., 2020). The MHQ 2016+ was intended for all participants who had completed the baseline assessment, regardless of whether they had participated in any subsequent assessments. Over 339,000 participants were invited via email, and by April 2025, more than 157,000 had completed the questionnaire.

In October 2022, the second online questionnaire on mental health, the Mental Well-being Questionnaire (MWB 2022+), was introduced (Davis, Coleman, et al., 2025). It included questions repeated from the MHQ 2016+, as well as new items addressing panic disorder, ED, general health and functioning, COVID-19, and resilience. This questionnaire was designed to be completed by all participants regardless of whether they had previously completed the MHQ 2016+. Over 329,000 participants were invited via email, and by April 2025, more than 170,000 had completed the MWB 2022+.

#### 2.2.2 Variable preprocessing

A common mistake when working with the UKB dataset is to assume that responses to similar or identical questions are coded uniformly across variables (Davis, Mirza, et al., 2025). This assumption can lead to three main issues: (a) erroneous inclusion of numeric codes when non-substantive responses are mistakenly treated as meaningful values, (b) variation in coding directionality, where higher values do not consistently reflect higher frequency, agreement, or intensity across variables, and (c) erroneous exclusion of meaningful responses when negative codes are mistakenly assumed to always represent non-substantive responses. To improve transparency and ensure consistent interpretation across variables, we systematically preprocessed variables to resolve these coding inconsistencies.

To comprehensively define missing values, we identified and excluded non-substantive responses based on their conceptual definitions. Specifically, responses whose descriptions matched patterns such as *Prefer not to answer* and *Do not know* were classified as missing. Notably, this text-based approach was necessary because non-substantive responses are not uniformly represented by negative codes across variables (e.g., 4 is used to indicate *Do not know* for a question about medication addiction, <u>Field 20551</u>), nor do all negative codes represent non-substantive responses. Beyond this, certain variables contained variable-specific codes that were treated as missing, including negative codes representing imprecise responses (e.g., −999 and −4 representing *Too many to count* for questions about unusual psychotic experiences), undocumented codes likely reflecting data entry artefacts (e.g., −1 for age when last took cannabis, <u>Field 20455</u>), and non-negative codes incompatible with the variable’s intended scale (e.g., 0 representing *Invalid weight* for body weight during illness, <u>Field 29125</u>; see **Table S2** for a full list). All such entries were subsequently excluded from analyses.

To ensure consistency in the interpretation of data coding, we identified variables in which the original coding was counterintuitive, that is, where higher numerical values corresponded to lower frequency or lower agreement (see **Table S3** for a full list of variables). For example, in response to questions about general happiness (e.g., <u>Field 4526</u>, <u>Field 20458</u>), the original coding assigned a value of 1 to indicate *Extremely happy* and 6 to indicate *Extremely unhappy*. To preserve interpretability and maintain consistent directionality across variables, we reverse-coded participants’ responses to these variables so that higher values reflect higher levels of the underlying construct.

To facilitate subsequent correlational analyses with IDPs, we transformed multi-response categorical variables into numeric variables by counting the number of options selected (see **Table S4** for a list of multi-response variables and the range of numeric coding). For example, for a question assessing leisure/social activities attended at least once per week (<u>Field 6160</u>), a participant who selected three activities was assigned a value of 3. Notably, a small subset of variables used negative codes to represent substantive response options, which were recoded to values that more accurately reflect their intended meaning prior to this transformation. For example, in a question assessing major life events in the past two years (<u>Field 6145</u>), the response option *None of the above* was originally coded as −7 and was recoded to 0. For a question assessing manic symptoms (<u>Field 6156</u>), the response option *All of the above* is mutually exclusive with the other four options representing individual symptoms. To preserve the meaning of this response, participants who selected it were assigned a value of 4.

#### 2.2.3 Repeated questions across assessment waves

We systematically identified questions that were repeated across different assessment waves. These included questions assessed repeatedly within the T/Screen 2009+ across all Assessment Centre visits, questions first introduced in the MHQ 2016+ and later repeated in the MWB 2022+, and questions spanning both the touchscreen and online questionnaires, the latter allowing responses to be tracked across up to six time points. This process was initially conducted by author XL and subsequently cross-checked by authors SK and LO.

#### 2.2.4 Overview of mental health phenotypes

Overall, we identified a total of 490 MHPs in the UKB. These comprised 48 phenotypes from the T/Screen 2009+ at the initial assessment visit, including 40 core items, 2 pilot items, and 6 derived measures. Only the core items were repeated, giving 40 phenotypes at each of the three subsequent Assessment Centre visits. The online questionnaires contributed a further 141 phenotypes from the MHQ 2016+ and 181 from the MWB 2022+. Together, these MHPs spanned 17 conceptual categories within three mental health domains: mental disorders, transdiagnostic features, and environmental factors. **Table 1** summarises the number of MHPs assessed across these domains and categories.

**Table 1.**
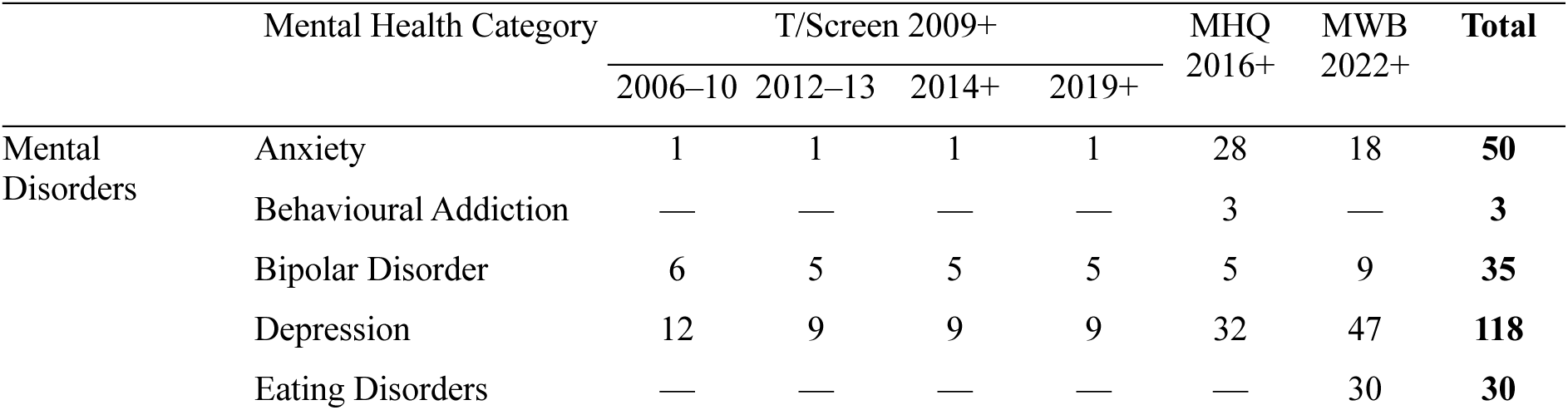
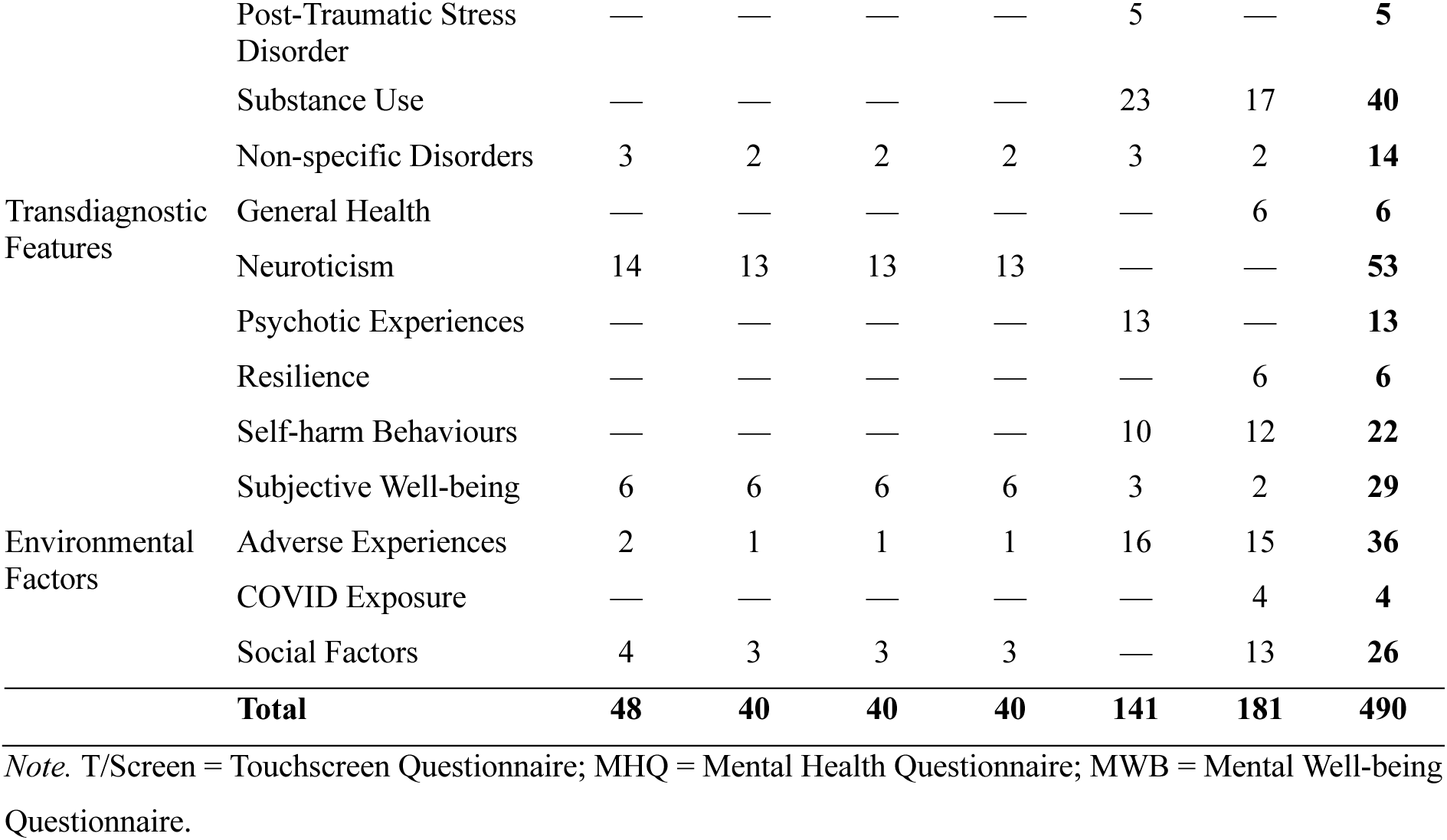
Number of phenotypes across mental health domains and assessment waves.

Among these phenotypes, 112 distinct questions were assessed at more than one time point. Of these, 29 questions from the T/Screen 2009+ were measured across up to four Assessment Centre visits, and 72 questions appeared in both the MHQ 2016+ and MWB 2022+, providing data at two time points. Notably, 10 questions assessing depressive symptoms and mood changes were included in both the touchscreen and online questionnaires, allowing responses to be tracked across up to six time points. In addition, one question from the T/Screen 2009+ corresponded to two questions in MWB 2022+, enabling responses to be followed across up to five time points (see **Table S5** for details).

### 2.3 Imaging derived phenotypes

#### 2.3.1 Data description

The UKB collects multimodal brain imaging data to characterize brain phenotypes relevant to a broad range of mental-health outcomes. The imaging protocol includes seven MRI modalities: T1-weighted MRI (T1), T2-weighted MRI (T2), susceptibility-weighted imaging (SWI), diffusion-weighted MRI (dMRI), task-based functional MRI (tfMRI), resting-state functional MRI (rfMRI), and ASL. Together, these modalities provide complementary information on anatomical structure, local tissue microstructure, and brain activity. Full details of the imaging acquisition protocol are described in Miller et al. (2016).

To enable large-scale analyses, the raw imaging data are processed through a fully automated pipeline that extracts quantitative summary measures from the MRI scans. These measures are referred to as IDPs and represent numerical indicators of brain structure or function, such as regional brain volumes, cortical thickness, white matter microstructure metrics, and functional connectivity estimates (Alfaro-Almagro et al., 2018; Miller et al., 2016).

T1 offers the most detailed depiction of brain anatomy, including clear delineation of grey and white matter and major anatomical landmarks, and is parcellated into regional measures using several standard atlases, including the Desikan-Killiany atlas (Desikan et al., 2006). Cortical and subcortical *volumes* derived from T1 are sensitive markers of tissue loss, capturing global atrophy and focal reductions. Cortical morphometry IDPs, including *cortical thickness* and *surface area*, offer additional insight into cortical atrophy and are clinically relevant. Intensity-based IDPs, including *mean T1 intensity* and *grey/white matter intensity contrast*, reflect regional tissue properties and are sensitive to variation across aging (Salat et al., 2009) and in psychiatric conditions (Jorgensen et al., 2016; Kong et al., 2015).

T2 is particularly valuable for detecting focal white matter lesions, which appear as hyperintensities on the images. The *total volume of white matter hyperintensities* is available as an IDP, offering a quantitative marker of lesion burden that are increasingly used in studies of mental-health conditions (Wang et al., 2014; Zhou et al., 2024).

SWI provides measures of magnetic susceptibility in deep grey-matter structures. Related IDPs in the UKB include *T2* signal decay times* and *quantitative susceptibility mapping* (QSM) values for subcortical nuclei, both of which are sensitive to tissue iron deposition.

dMRI is sensitive to the random motion of water molecules, which is affected by tissue microstructure. Since diffusion is typically greater along the length of axons than across them, dMRI can be used to infer the organisation of white matter tracts. In the UKB, two complementary models are fit to the diffusion signal to derive microstructural metrics. The diffusion-tensor model produces *fractional anisotropy* (FA), which quantifies the degree to which diffusion is directionally constrained, *mean diffusivity* (MD), reflecting the overall magnitude of diffusion averaged across directions, and *mode of anisotropy* (MO), which describes the geometric shape of the diffusion tensor. The three *principal diffusivities* (L1, L2 and L3) correspond to the eigenvalues of the diffusion tensor and describe the magnitude and represent the magnitude of diffusion along the three principal axes. The neurite orientation dispersion and density imaging (NODDI) model provides *intracellular volume fraction* (ICVF), *orientation dispersion* (OD) and *isotropic volume fraction* (ISOVF), which estimate neurite density, fibre complexity and free-water content, respectively. Whole-brain structural connectomes computed from the dMRI data are available as a returned dataset (<u>Fields</u> <u>31020</u>–<u>31026</u>; Mansour et al., 2023), but were contributed by an individual research group rather than generated by the standard pipeline (Alfaro-Almagro et al., 2018), and are therefore not included among the IDPs analysed here.

Functional MRI measures brain activity indirectly through fluctuations in blood oxygenation and flow that accompany changes in neural metabolic demand. tfMRI assesses blood-oxygen-level-dependent (BOLD) responses during an explicit task. In the UKB protocol, participants completed a faces-shapes emotion-matching paradigm designed to elicit activation in perceptual and emotion-processing regions (Hariri et al., 2002). IDPs derived from this acquisition include the *BOLD effect* and the *z-statistic* for specific task contrasts, extracted within regions defined from group-averaged activation maps. These IDPs quantify the strength of task-evoked activation and the associated statistical evidence in predefined functional regions.

rfMRI measures spontaneous BOLD fluctuations in the absence of a task, providing a way to study intrinsic brain networks. Alterations in these networks have been widely reported across mental illnesses (Sheline et al., 2010; Woodward & Cascio, 2015). In the UKB, group-level independent component analysis (ICA) is used to decompose the resting-state signal into a set of functionally defined components, each representing a node in the intrinsic network. Two main types of IDPs are derived from these nodes. *Component amplitudes* quantify how strongly each node fluctuates over time, providing a measure of node-level activity. *Functional connectivity* IDPs capture the strength of associations between pairs of nodes using full or partial correlations, summarising how different parts of the network communicate.

ASL provides a non-invasive measure of cerebral perfusion by magnetically labelling inflowing arterial blood as an endogenous tracer. The related IDPs have been available in the UKB since February 2023 and include regional measures of *cerebral blood flow* (CBF) and *arterial transit time* (ATT), reported separately across cortical regions and vascular territories. CBF quantifies tissue perfusion, whereas ATT reflects the time required for labelled blood to reach the tissue, providing information about vascular delivery and hemodynamic efficiency. The ASL measures are widely used to examine alterations in cerebral perfusion in neurological and psychiatric disorders (Haller et al., 2016).

#### 2.3.2 Variable preprocessing

rfMRI IDPs in UKB are stored as *bulk* or compound variables, where each data field contains a text file holding all values for each measure for a given subject. Individual IDPs therefore need to be extracted from these files. For component amplitudes, the bulk text file contains a row vector whose length matches the number of ICA components retained for a specific processing pipeline. For connectivity IDPs, UKB provides only the unique pairwise connections, which correspond to the upper-triangular entries of the symmetric connectivity matrix.

Since the rfMRI analyses are performed at several parcellation dimensionalities, which directly determine how many IDPs can be extracted, we also briefly summarise the details here. In the volumetric pipeline, ICA is performed at 25 and 100 dimensions, with artifactual components removed before defining the final set of nodes, resulting in 21 and 55 usable nodes. The surface-based pipeline, which has been available in UKB since March 2024, performs ICA at 25 and 50 dimensions and retains all components.

#### 2.3.3 Overview of imaging derived phenotypes

As of April 2025, 8,815 distinct IDPs are available for each imaging visit, giving a total of 17,630 IDPs. **Table 2** summarises the number of IDPs available across imaging modalities and categories of brain measures.

**Table 2.**
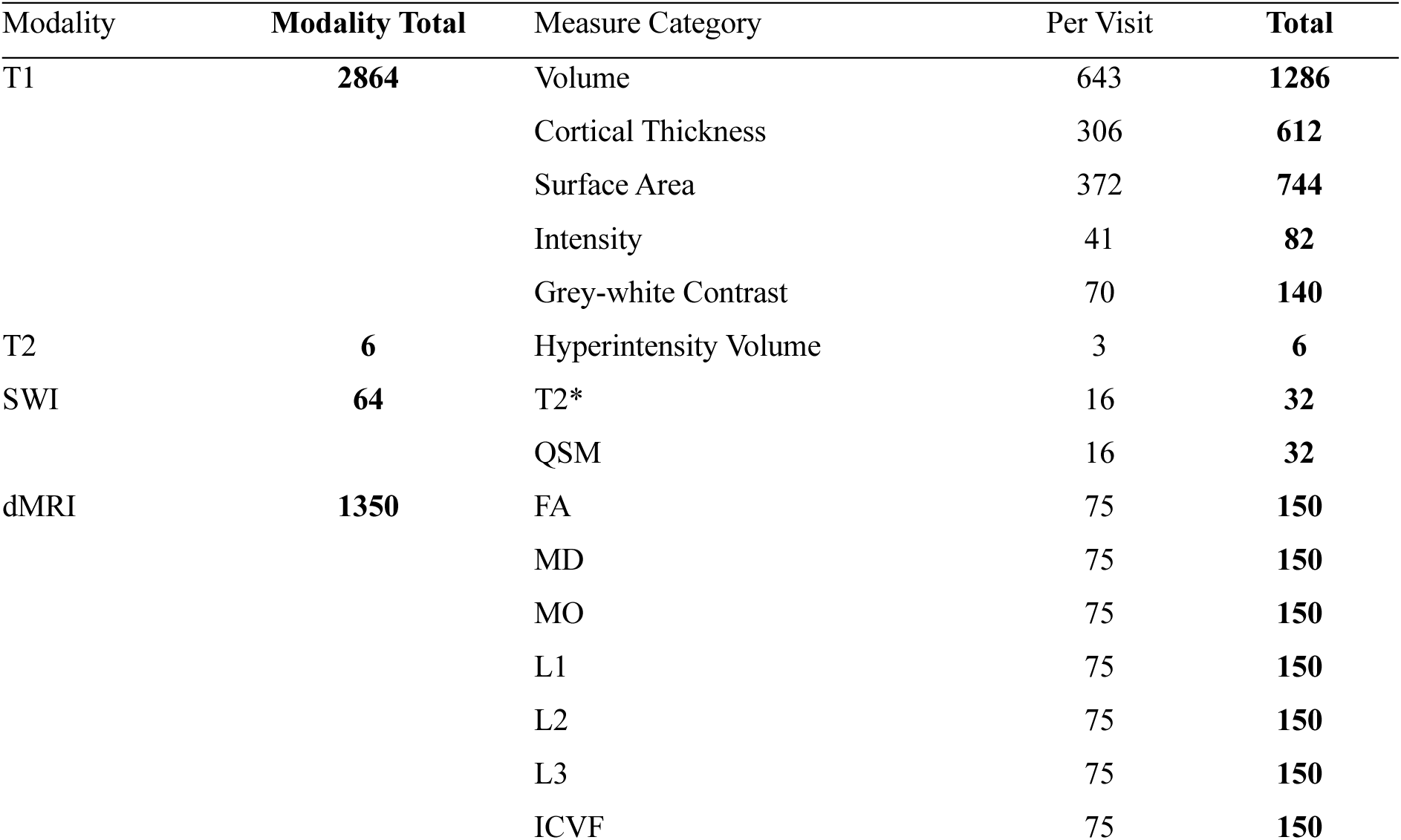
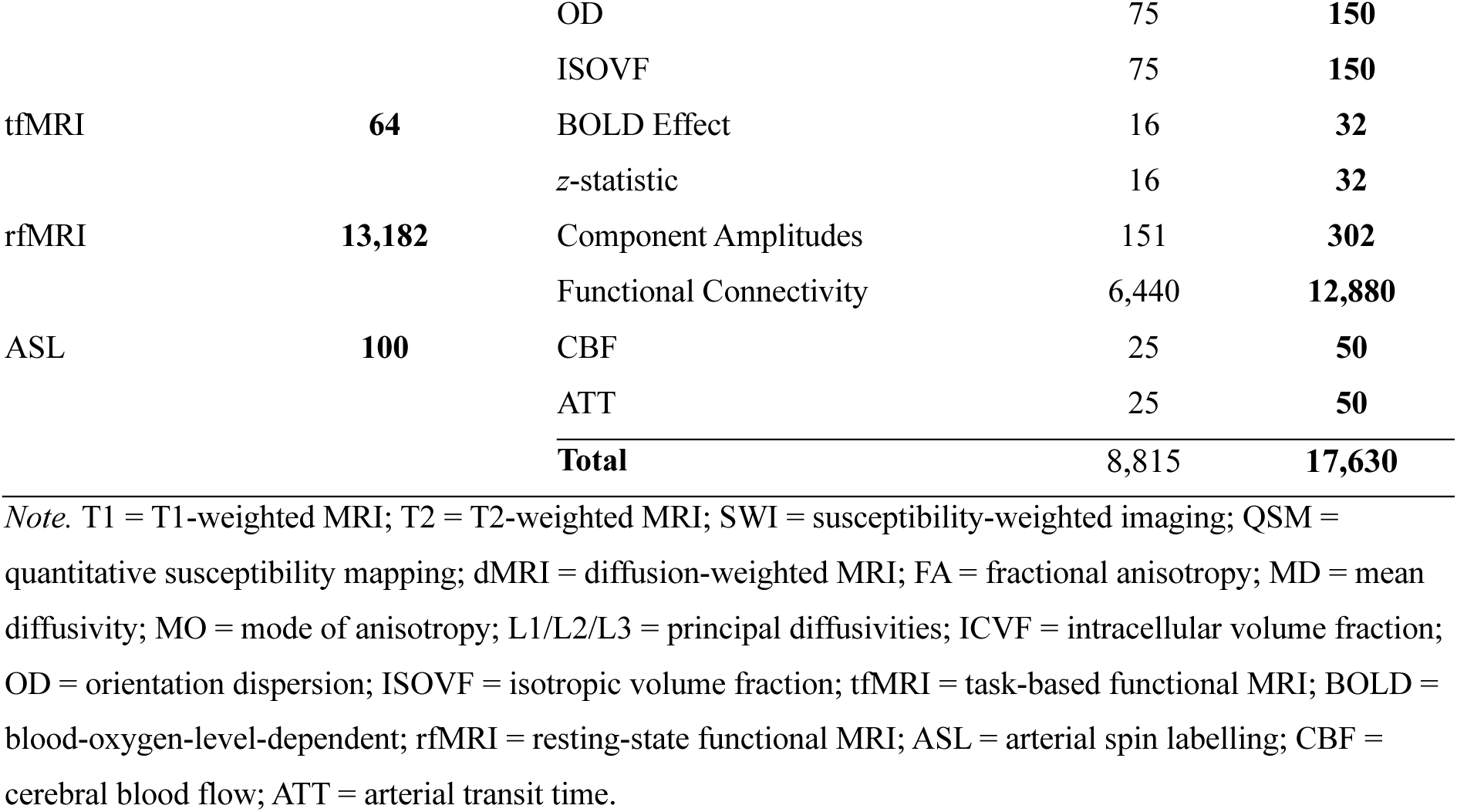
Number of imaging derived phenotypes across imaging modalities.

### 2.4 Brain-mental health association analysis

#### 2.4.1 Cohort and variable selection

For the brain-mental health association analyses, we used a sub-cohort of participants who completed both the initial and repeat imaging visits in 2014+ and 2019+, as well as both the MHQ 2016+ and MWB 2022+ online questionnaires. Because the touchscreen questionnaire was administered at every Assessment Centre visit, we restricted the touchscreen measures to the two imaging visits, which are the only occasions at which mental health and imaging data were collected together. This left 402 of the 490 identified MHPs, excluding the 88 assessed at the initial and first repeat assessment visits.

To ensure adequate variability and data completeness for subsequent analyses, we evaluated each variable against exclusion criteria established in previous population studies (Miller et al., 2016; Smith et al., 2015). Variables were excluded if they lacked any meaningful variance in the defined cohort (i.e., all participants provided identical responses), if responses were heavily dominated by a single option (i.e., more than 95% of participants provided the same response), or if the proportion of participants with valid responses was insufficient for reliable estimation (i.e., fewer than 10% of participants provided non-missing data). Of the 402 MHPs, none lacked variance entirely, 69 were excluded for insufficient valid responses (3 from the T/Screen 2009+, 27 from the MHQ 2016+, and 39 from the MWB 2022+) and 24 for a dominant response (2, 11, and 11, respectively), yielding a final set of 309 MHPs. All phenotypes in two categories, behavioural addiction and psychotic experiences, were excluded at this stage, so these categories are not represented in the association analyses. Most were administered only to participants who endorsed a preceding screening question and were excluded for insufficient valid responses (0.3% to 5.0% valid); the screening questions themselves were excluded for a dominant response, having been answered negatively by 97.2% to 99.4% of participants. Among the touchscreen phenotypes, one was excluded at the initial imaging visit and four at the repeat, leaving 39 and 36 respectively, with the 36 available at both visits. The same criteria excluded the 50 ASL IDPs from the initial imaging visit, since ASL was introduced late in the protocol and had not been acquired for most participants at that visit, giving a final set of 17,580 IDPs.

To validate the replicability of the associations, the dataset was then randomly divided into discovery (*N* = 4,207) and validation (*N* = 4,208) subsets. Characteristics at each visit were compared between subsets using independent-samples *t*-tests for continuous variables, chi-squared tests for categorical variables, and standardised Mann-Whitney U tests for date-related variables.

#### 2.4.2 Confound variables

Controlling for confounds is essential for identifying true associations in large-scale studies, where demographic and technical factors can introduce substantial variance in both imaging and mental health measures (Alfaro-Almagro et al., 2021). At the same time, variables should not be treated as confounds unless they plausibly cause systematic bias in the measurement, as inappropriate adjustment can remove true effects or create spurious ones. In this project, we included confounds only when they met this criterion. Unlike previous analyses that applied a uniform confound set across all variables (Dutt et al., 2022), we identified separate sets of confound for IDPs, MHPs from the touchscreen questionnaire during imaging visits, and MHPs from online questionnaires.

For all IDPs, we used the “simple” set of confounds described in Alfaro-Almagro et al. (2021), which was shown to explain 4.4% of variance in the UKB imaging variables on average. It included scanning site, age at assessment centre visit, age², sex, age × sex interaction, head size, head motion in rfMRI and tfMRI, scan date, and scan date². Although UKB uses identical scanner hardware and software across all imaging sites, we included acquisition site as a confound to account for other subtle differences (Alfaro-Almagro et al., 2021). Quadratic terms for age and scan date were included to model potential nonlinear relationships between these variables and the IDPs. Age-related changes in brain structure often follow nonlinear trajectories across the adult lifespan (Bethlehem et al., 2022).

Similarly, date² captures potential nonlinear temporal trends in data collection. We also included an age × sex interaction term to capture differential age-related trajectories between males and females (Bethlehem et al., 2022).

For MHPs acquired during imaging visits, we also included scanning site, age, age², sex, age × sex interaction, scan date, and scan date² as confounds. We retained scanning site since visit location could plausibly influence self-report responses through factors like local participant demographics, travel burden to the imaging centre, or site-specific assessment procedures.

These confounds were generated separately for each imaging visit and each subject subset. Continuous confounds (age, head size, motion, scan date) were first normalized globally across all subjects, then duplicated by site to allow site-specific effects while keeping these variables independent of the site indicator variables (Alfaro-Almagro et al., 2021).

Categorical confound (sex) was similarly coded separately for each site. Quadratic and interaction terms were then computed from these site-specific variables and orthogonalized to the linear confounds.

For MHPs from online questionnaires, we included age at questionnaire completion, age², sex, age × sex, questionnaire completion date, and date² as confounds. Since online questionnaires were completed independently of imaging visits, all confounds were normalized globally for each subject subset. Quadratic and interaction terms were computed from the normalized variables and orthogonalized to all linear confounds.

#### 2.4.3 Pearson’s correlations

To examine between-subject brain-mental health associations, we computed Pearson’s correlations between MHPs and IDPs. Both phenotype datasets were first rank-based inverse normal transformed to approximate Gaussian distributions, then deconfounded using linear regression. The resulting residuals were then used for the correlation analyses, with missing data being handled by pairwise deletion. All analyses were conducted separately for the discovery and validation subsets.

The correlations were then summarised using a common set of metrics throughout. *Median |r|* and the 99^th^ percentile of |*r*|, reported as the *top 1% |r|*, describe the centre and upper tail of the effect-size distribution. Replicability was primarily assessed by the *replication rate*, defined as the proportion of associations reaching *p* < 0.05 in discovery that also reached *p* < 0.05 in validation, irrespective of the sign of the correlation. All metrics are computed within whichever set of associations is being described, whether the full matrix or a domain, category, modality, or category-modality combination. FDR correction was likewise applied within each such set rather than across the full matrix. Where replication rate is plotted as a function of effect size, associations are grouped into equal-width bins of |*r*| and the rate is computed within each bin.

#### 2.4.4 Repeated measures correlations

For the repeated measures analysis, we identified confounds as variables that change within subjects across imaging visits. For IDPs, time-varying confounds included head size, head motion in rfMRI and tfMRI, and scan date. For MHPs collected during imaging visits, only scan date was treated as a time-varying confound. Age was not treated as a confound in the repeated-measures correlation analyses because age-related change is part of the processes of interest. The resulting effect sizes reflect within-subject associations that include coordinated age-related changes, providing benchmarks appropriate for researchers designing longitudinal studies of brain-mental health relationships.

To examine within-subject associations across time, we computed repeated measures correlations between MHPs and IDPs using the *rmcorr* R package (Bakdash & Marusich, 2017). Because repeated measures correlations require a measurement at each visit, these analyses were restricted to the 8,765 IDPs and 36 touchscreen MHPs available at both imaging visits. Both MHPs and IDPs, along with their respective time-varying confounds, were rank-based inverse normal transformed separately at each timepoint. Phenotype data were then deconfounded for these time-varying confounds prior to analysis. Time-invariant variables, including sex and scanning site, were not explicitly deconfounded because the model inherently accounts for stable between-subject differences by estimating a common within-subject association across timepoints. All correlations were performed separately in the discovery and validation subsets.

## 3 Results

### 3.1 Cohort description

The final cohort for analysis comprised 8,415 participants (53.7% female) with a mean age of 62.84 years (*SD* = 7.36, range = 45–81) at the initial imaging visit. Participants were recruited from three assessment sites: Cheadle (43.0%), Newcastle (53.0%), and Reading (4.0%). The initial imaging visit was conducted between May 2014 and May 2022, and the repeat imaging visit between May 2019 and October 2024. The interval between the two imaging visits had a median of 4.99 years (IQR = 2.31–5.74, range = 1.09–10.33). The MHQ 2016+ and MWB 2022+ online questionnaires were completed within much narrower windows, spanning July 2016 to July 2017 and October 2022 to August 2023 respectively, with the majority of responses received within the first two months of each questionnaire being sent out.

Discovery and validation subsets were comparable across all cohort characteristics (all *p* > 0.05; **Table S6**).

### 3.2 Overall effects

Across the full dataset, brain-mental health associations in UKB were predominantly very small. Among more than 5.4 million correlations between 309 MHPs and 17,580 IDPs, the median absolute effect size was |*r*| = 0.018, the top 1% exceeded |*r*| = 0.108, and the largest reached |*r*| = 0.430 in the discovery subset (**Figure 1a**). The signed distribution was approximately bell-shaped and centred near zero, with no systematic tendency toward positive or negative associations. The validation subset showed the same distribution (median |*r*| = 0.018; top 1% |*r*| = 0.106).

**Figure 1.**
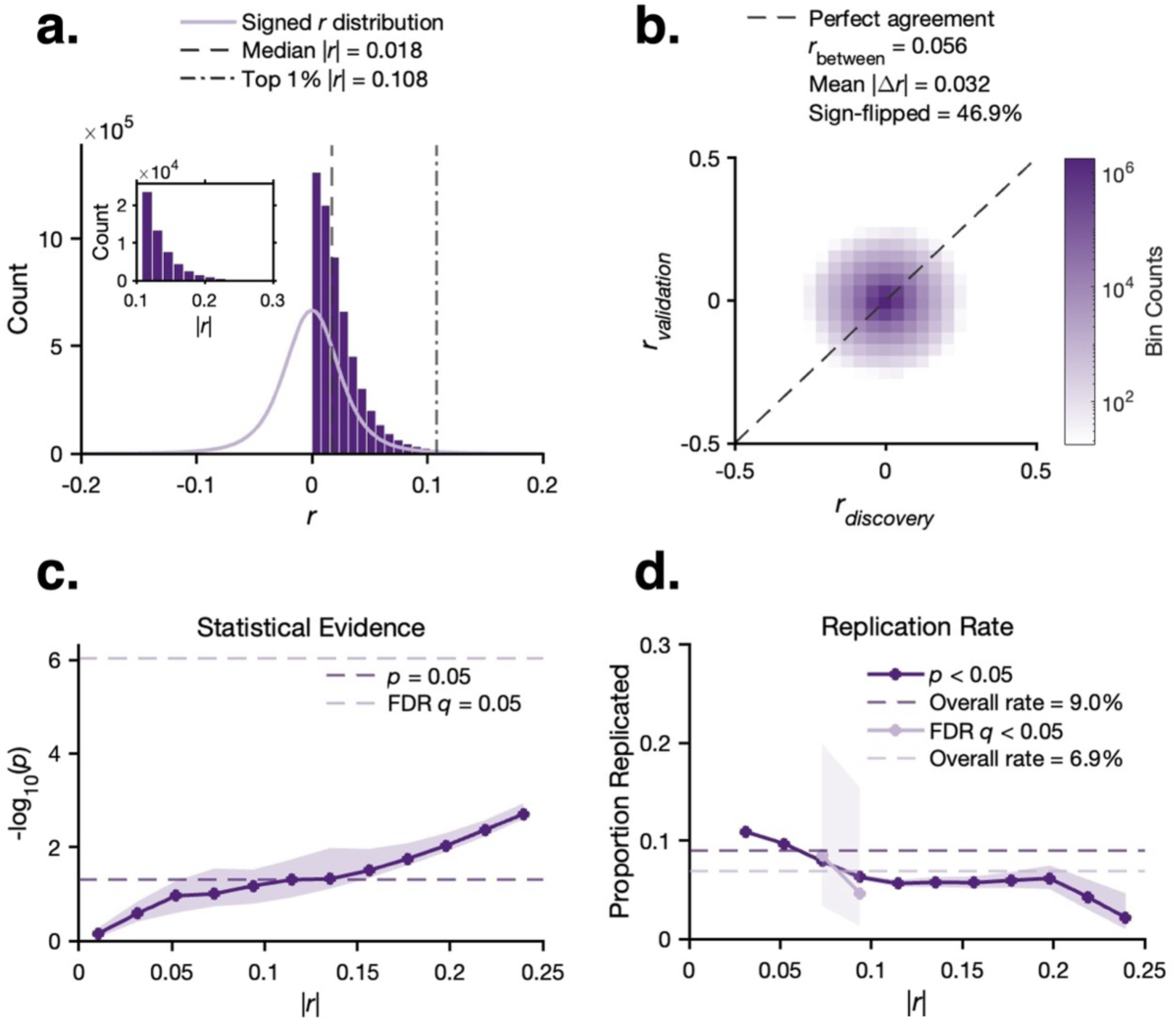
Effect size and replicability of brain-mental health associations. Pearson correlations between 17,580 IDPs and 309 MHPs were computed in discovery (*N* = 4,207) and validation (*N* = 4,208) subsets, yielding approximately 5.4 million pairwise tests per subset. **(a)** Distributions of absolute correlation coefficients (|*r*|; dark purple bars) in the discovery subset, with the kernel density estimate of the signed correlation distribution overlaid (light purple curve). *Inset:* zoomed histogram of the upper tail of |*r*|. **(b)** Two-dimensional density scatter plot of discovery against validation correlation coefficients. The dashed diagonal marks perfect between-subset agreement; colour represents bin counts on a logarithmic scale. **(c)** Statistical evidence as a function of effect size, shown as the median −log_10_(*p*) within equal-width bins of |*r*|. The shaded ribbon indicates the interquartile range across associations in each bin. **(d)** Replication rate as a function of effect size, shown at each of two significance threshold, *p* < 0.05 and FDR-corrected *q* < 0.05. Lines show the proportion of discovery-significant associations that were also significant in validation within each |*r*| bin. Shaded ribbons show 95% Wilson confidence intervals; wider ribbons reflect bins with fewer significant associations and correspondingly less stable estimates. Bins with fewer than 10 significant associations are suppressed.

6.3% of discovery correlations reached *p* < 0.05 (6.4% in validation), with significance becoming attainable once |*r*| exceeded roughly 0.07 (**Figure 1c**). Replicability across the two subsets was considerably poorer. Of the discovery associations reaching *p* < 0.05, 9.0% were also significant in validation, and of the 101 surviving FDR correction, 6.9% remained significant. The effect-size estimates showed almost no agreement between subsets (*r*_between_ = 0.056, mean |Δ*r*| = 0.032), reflected visually in the density scatter plot as a roughly circular cloud centred on the origin rather than an elongation along the diagonal (**Figure 1b**), and 46.9% of associations flipped sign across subsets, close to the 50% expected when the true effects are near zero and the sign is determined mainly by sampling noise. Replication itself inversely depended on effect size, falling from 11% at |*r*| = 0.03 to 2% beyond |*r*| = 0.20 (**Figure 1d**).

### 3.3 Mental health domain- and category-specific effects

Mental disorders showed the largest effects of the three domains (median |*r*| = 0.020; top 1% |*r*| = 0.118) but replicated in only 7.8% of cases. Transdiagnostic features showed smaller effects (median |*r*| = 0.015; top 1% |*r*| = 0.074) yet replicated most often (12.3%), and environmental factors, though comparable in magnitude (median |*r*| = 0.015; top 1% |*r*| = 0.072), replicated in 8.9% of cases. Effect size and replicability therefore diverged at the domain level, with the largest associations found where they were least likely to replicate (**Table S7**).

#### 3.3.1 Mental disorders

ED showed the largest effects of any mental disorder category (median |*r*| = 0.033; top 1% |*r*| = 0.177) but replicated in only 6.0% of cases (**Figure 2a**), and BD, with the second-largest effects, replicated least within the domain (5.0%). Substance use showed more modest effects but the highest replication (14.5%). The two largest categories, depression and anxiety, sat near the domain average in both magnitude and replication (6.7% and 7.0% of associations replicating), as did post-traumatic stress disorder (PTSD), the smallest category. Non-specific diagnoses replicated somewhat more often (9.4%).

**Figure 2.**
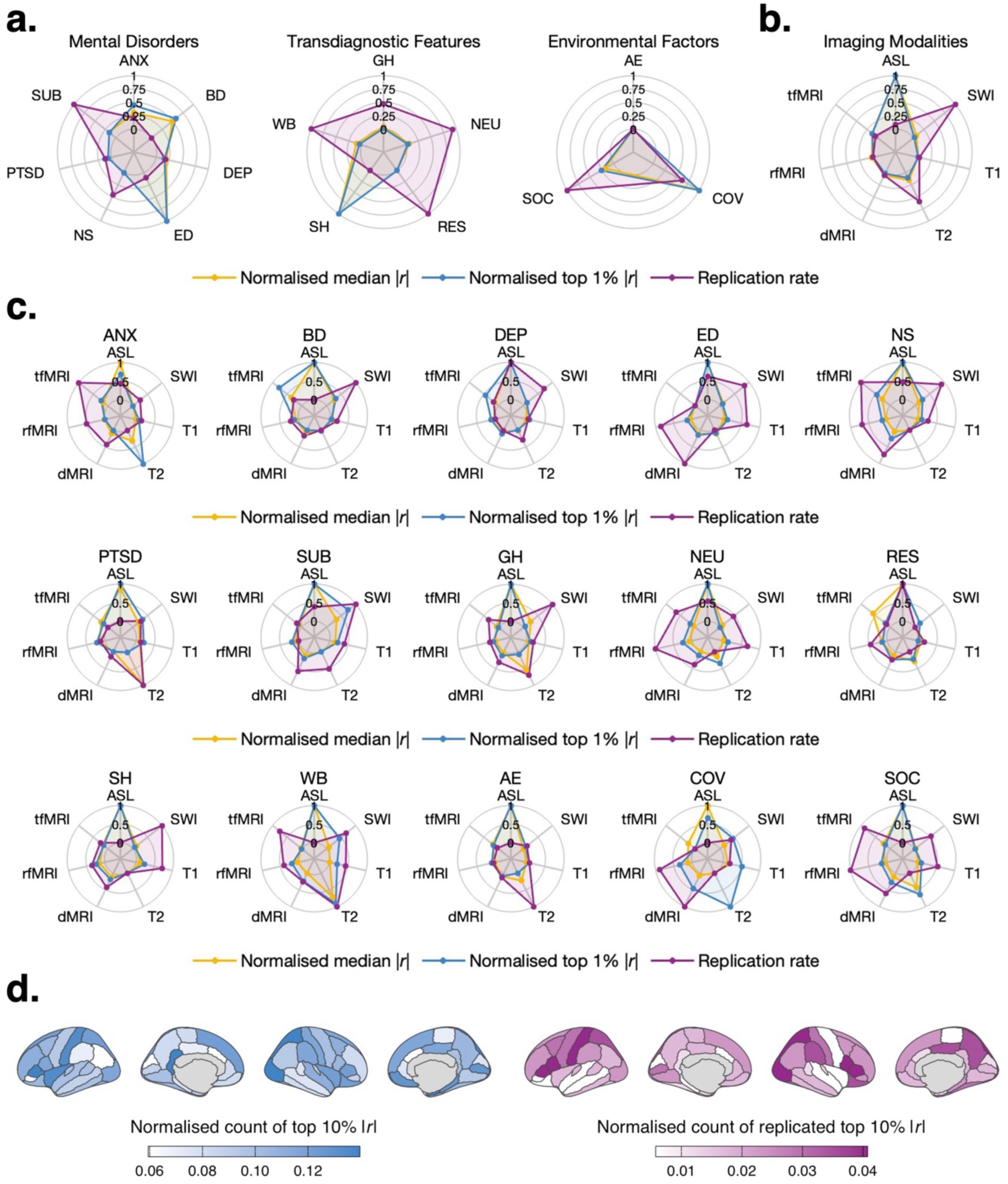
Effect size and replicability of brain-mental health associations across mental health domains and categories and imaging modalities. In all radar plots, median |*r*|, top 1% |*r*|, and replication rate are min-max normalised across the spokes of each plot, so plotted values reflect relative magnitude. **(a)** Domain-level plots, one per mental health domain, with one spoke per mental health category. **(b)** Modality-level plot, with one spoke per imaging modality. **(c)** Category-by-modality plots, one per mental health category, with one spoke per imaging modality; categories are grouped by domain, read *left* to *right* and *top* to *bottom*. **(d)** Associations between depression phenotypes and regional mean cortical thickness, with regions defined by the Desikan-Killiany atlas and shown as lateral and medial views of both hemispheres. The magnitude map (*left*) colours each region by the share of its correlations whose |*r*| falls in the strongest 10% across the cortex in the discovery subset; the replication map (*right*) shows the share ranking in the strongest 10% in both subsets. Colour is scaled to the 5^th^ to 95^th^ percentile of the plotted values in both maps and regions not covered by the atlas are shown in grey. ANX = anxiety; BD = bipolar disorder; DEP = depression; ED = eating disorders; NS = non-specific diagnoses; SUB = substance use; GH = general health; NEU = neuroticism; RES = resilience; SH = self-harm thoughts and behaviours; WB = subjective well-being; AE = adverse experiences; COV = COVID exposure; SOC = social factors; ASL = arterial spin labelling; SWI = susceptibility-weighted imaging; T1 = T1-weighted MRI; T2 = T2-weighted MRI; dMRI = diffusion-weighted MRI; rfMRI = resting-state functional MRI; tfMRI = task-based functional MRI.

To identify which imaging modalities contributed most strongly to associations within each mental health category, we examined effect size and replicability for each category-modality combination (**Figure 2c**; **Table S9**). ASL produced the largest within-category effects in most mental disorder categories, with effects exceeding those of any other modality on both median and top 1% |*r*| for BD, depression, ED, and substance use. Associations with ASL IDPs were computed on a relatively smaller subsample than associations with other modalities, so the ASL effects should be read as preliminary. Setting ASL aside, tfMRI was the strongest modality for BD (median |*r*| = 0.029, top 1% |*r*| = 0.186) and depression (median |*r*| = 0.021, top 1% |*r*| = 0.125). For ED, dMRI was the next strongest modality (median |*r*| = 0.037, top 1% |*r*| = 0.180), with the highest replication rate within the category (7.1%). For substance use, SWI was the next strongest modality (median |*r*| = 0.024, top 1% |*r*| = 0.124) and replicated in 39.3% of cases. T2 led on top 1% |*r*| for anxiety (0.204) and on median |*r*| for PTSD (0.036), though the latter rests on 30 associations and replicated in 50% of them, against 4.8% for anxiety.

Beyond the modality level, many IDPs are defined for specific brain regions, allowing us to examine the regional distribution of brain-mental health associations. To show what that resolution yields, **Figure 2d** maps how cortical thickness was associated with depression phenotypes across the cortex. The magnitude map (*left*) shades each region by how often its effects ranked among the strongest 10% across the whole cortex, and the replication map (*right*) by how often those associations also held in the validation subset. Several regions stood out on both, including the bilateral pars triangularis and opercularis, the left postcentral cortex, and the right superior parietal and lateral occipital cortex.

#### 3.3.2 Transdiagnostic features

Self-harm thoughts and behaviours stood out for having the largest median and top 1% effects among the transdiagnostic features (median |*r*| = 0.019; top 1% |*r*| = 0.130) yet showed the lowest replication rate (6.3%; **Figure 2a**). Resilience, subjective well-being, and neuroticism showed smaller effects but much higher replication rates (13.6%, 13.5%, and 13.0%, respectively). General health matched these three in magnitude but showed a lower replication rate (9.7%).

By modality, ASL again produced the largest effects for most features (**Figure 2c**). This was most pronounced for self-harm thoughts and behaviours, where ASL exceeded every other modality by a factor of three or more (median |*r*| = 0.070; top 1% |*r*| = 0.280), and no clear second modality emerged: tfMRI was marginally higher on median |*r*| (0.021), while rfMRI, T1, and dMRI clustered on top 1% |*r*| (0.120–0.130). Neuroticism followed the same pattern at smaller magnitude, with ASL leading on median |*r*| (0.025), the remaining modalities undifferentiated (0.014–0.016), and replication highest for rfMRI (14.2%). For resilience, ASL led on both magnitude (median |*r*| = 0.030, top 1% |*r*| = 0.089) and replication (33.3%), with tfMRI a clear second on median |*r*| (0.023). For subjective well-being and general health, T2 was the strongest modality after ASL (median |*r*| = 0.029 and 0.026) and replicated in 33.3% and 37.5% of cases.

#### 3.3.3 Environmental factors

COVID-19 exposure showed the largest effects (median |*r*| = 0.019; top 1% |*r*| = 0.111) and a relatively low replication rate (9.8%) among the environmental factors (**Figure 2a**). Social factors showed slightly smaller effects but the highest replicability (11.5%). Adverse experiences showed similar effect sizes but the lowest replication (7.0%).

By modality, ASL again produced the largest median effects across all three categories (median |*r*| = 0.033, 0.039, and 0.033 for adverse experiences, COVID exposure, and social factors; **Figure 2c**). T2 was a recurring second on median |*r*| for adverse experiences (0.019) and social factors (0.022). For COVID exposure, T2 gave the highest top 1% |*r*| (0.156), though with only 24 associations this reflects little more than the single largest correlation. T2 was the most replicable modality for adverse experiences (43.8%), whereas rfMRI was most replicable for social factors (12.7%) and dMRI for COVID exposure (13.0%).

### 3.4 Imaging modality-specific effects

Effects were broadly similar across modalities, with the exception of ASL, but replication varied more than twofold (**Figure 2b**; **Table S8**). ASL showed the largest effects of any modality (median |*r*| = 0.038; top 1% |*r*| = 0.232), with moderate replication (9.7%). T1 showed relatively small effects (median |*r*| = 0.017; top 1% |*r*| = 0.108) and moderate replication (9.1%). T2 showed somewhat larger effects (median |*r*| = 0.021; top 1% |*r*| = 0.120) and higher replication (14.6%). SWI showed comparable effects (median |*r*| = 0.019; top 1% |*r*| = 0.109) but the highest replication of any modality (18.4%). dMRI was similar in effect magnitude (median |*r*| = 0.018; top 1% |*r*| = 0.109) with moderate replication (9.4%). rfMRI (median |*r*| = 0.018; top 1% |*r*| = 0.107) and tfMRI (median |*r*| = 0.020; top 1% |*r*| = 0.123) showed similarly small effects with moderate replication rates of 8.8% and 9.3%, respectively.

### 3.5 Longitudinal structure of brain-mental health relationships

The initial and repeat imaging visits (2014+ and 2019+) were the only assessments at which brain imaging and mental health measures were acquired on the same day. Having both measurements at two time points allows brain-mental health associations to be examined across time in two ways: how the associations behave across different temporal configurations of measurement, and whether brain and mental health change together within individuals. Both analyses used the 36 touchscreen MHPs common to both visits and the 8,765 IDPs available at each visit, in the same discovery and validation subsets as above.

#### 3.5.1 Between-subject associations across visits

Temporal separation between the brain and mental health measurements did not weaken their association. Because both the IDPs and the touchscreen MHPs were available at each visit, the two could be paired within a visit or across visits, giving two same-day and two cross-visit configurations. The *left* panel of **Figure 3a** plots the cumulative distribution of effect size for the two same-day configurations. Correlating initial-visit IDPs with initial-visit MHPs gave smaller effects and higher replication (median |*r*| = 0.013; top 1% |*r*| = 0.055; replication rate = 13.6%) than correlating repeat-visit IDPs with repeat-visit MHPs (median |*r*| = 0.019; top 1% |*r*| = 0.086; replication rate = 9.2%), with the former curve rising more steeply, so more of its associations sit at low |*r*|, and the latter extending further into the upper tail. This separation alone might suggest that associations strengthened across the roughly five years between visits. Setting these against the cross-visit configurations (**Figure 3a**, *right*) tells a different story. Curves built on brain measures from a given visit nearly overlie each other across the two panels: pairing initial-visit IDPs with initial- or repeat-visit MHPs made almost no difference (median |*r*| = 0.013; top 1% |*r*| = 0.055 and 0.057; replication rate = 13.6% and 13.1%), and the same held for repeat-visit IDPs (median |*r*| = 0.019; top 1% |*r*| = 0.086 and 0.081; replication rate = 9.2% and 7.6%). The separation therefore tracked which visit the brain measures came from, not the temporal gap between measurements, and held regardless of when the mental health measure was taken or which was assessed first. In the discovery subset, few associations survived FDR correction in any configuration, and essentially none did in validation (**Table S10**).

**Figure 3.**
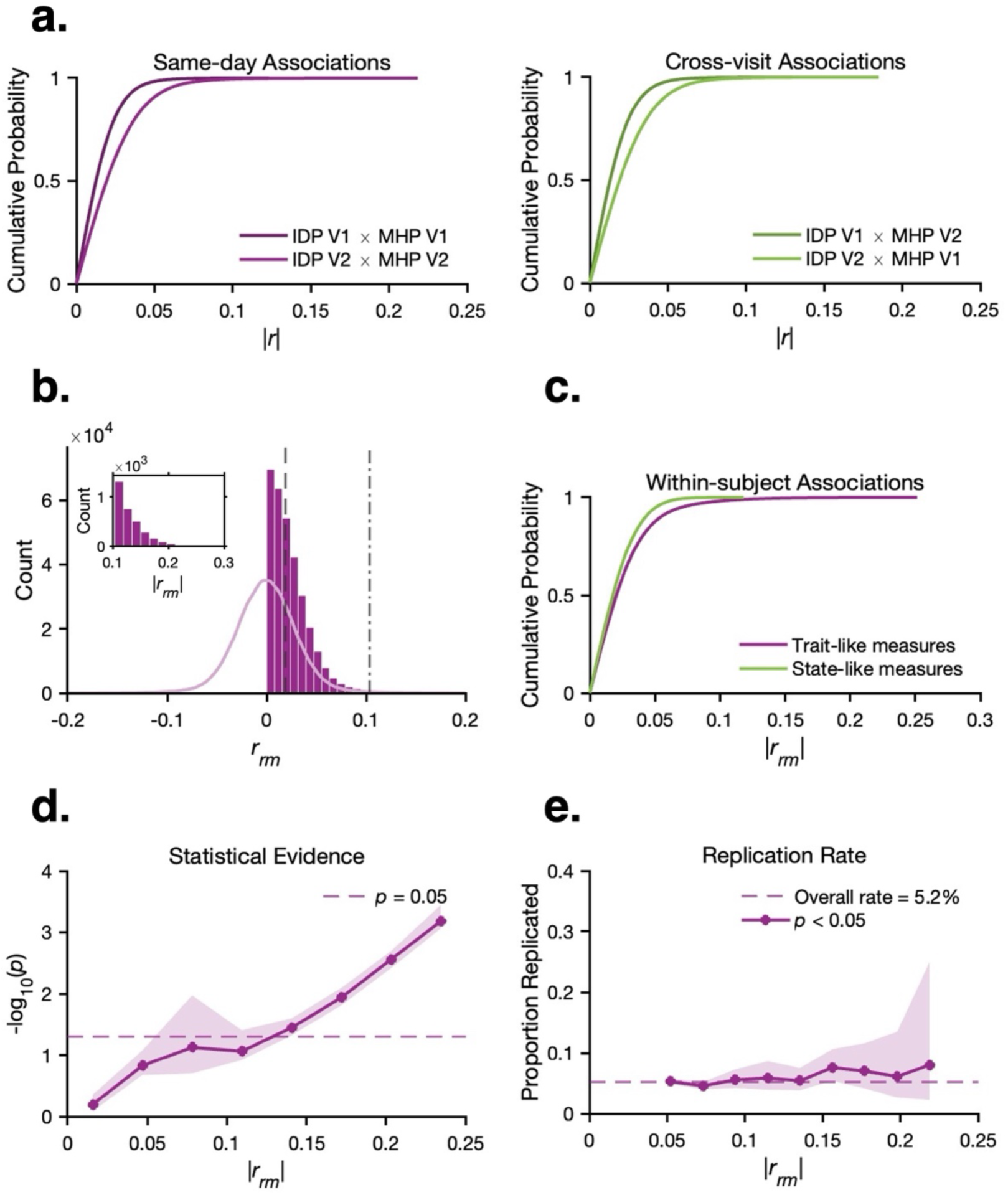
Same-day, cross-visit, and within-subject brain-mental health associations across the two imaging visits. Pearson’s and repeated measures correlations between the 36 touchscreen MHPs and 8,765 IDPs available at both imaging visits were computed in the discovery and validation subsets. **(a)** Between-subject associations as empirical cumulative distributions of |*r*| in the discovery subset, for the two same-day configurations (*left*) and the two cross-visit configurations (*right*), where V1 and V2 denote the initial and repeat imaging visits. **(b)** Distribution of signed within-subject repeated-measures correlations (*r_rm_*) in the discovery subset, shown as a histogram with the kernel density estimate of the signed distribution overlaid. *Inset*: the upper tail of |*r_rm_*|. Dashed and dash-dot lines mark the median (|*r_rm_*| = 0.019) and top 1% (|*r_rm_*| = 0.104). **(c)** Empirical cumulative distributions of |*r_rm_*| for trait-like and state-like MHPs in discovery. **(d)** Statistical evidence for the repeated-measures correlations, shown as the median −log10(*p*) within bins of |*r_rm_*| in discovery, with the shaded ribbon showing the interquartile range across associations in each bin; the dashed line marks *p* = 0.05. **(e)** Replication rate of the repeated-measures correlations, shown as the proportion of discovery-significant associations (*p* < 0.05) also significant in validation within bins of |*r_rm_*|; shaded ribbons show 95% Wilson confidence intervals, and the dashed line marks the overall rate. Bins with fewer than 10 significant associations are suppressed.

#### 3.5.2 Within-subject change across visits

Repeated-measures correlations showed that, within participants, changes in brain measures across the two imaging visits showed little association with changes in mental health measures. Since age was not partialled out as a confound for this analysis, any age-related change shared by brain and mental health measures would inflate these correlations. That they remained close to zero therefore indicates the null is not a consequence of removing age-related variance. In the discovery subset, the correlations centred near zero (median |*r_rm_*| = 0.019; top 1% |*r_rm_*| = 0.104; maximum |*r_rm_*| = 0.252; **Figure 3b**). The validation subset was near-identical (median |*r_rm_*| = 0.019; top 1% |*r_rm_*| = 0.097). Associations became significant only above |*r_rm_*| ≈ 0.12 (**Figure 3d**). About 5% of correlations reached significance in each subset, and none survived FDR correction (**Table S11**).

Replicability was low. Agreement between the two subsets was negligible (*r*_between_ = −0.004), and 49.6% of associations changed direction between them. The mean absolute difference between the effect-size estimates was small (|Δ*r_rm_*|= 0.033, *SD* = 0.030). Only 5.2% of discovery-significant associations were also significant in validation, and the larger within-subject associations replicated no more often than the smaller ones, with the replication rate in **Figure 3e** staying flat near the overall average as effect size increases.

To test whether the result depended on how much a measure can change within a person, we compared associations for a set of trait-like measures (*n* = 22), such as lifetime symptoms, diagnoses, and personality traits, with those for the state-like measures of recent symptoms, mood, and circumstances (*n* = 14). Both distributions concentrated near zero, although the trait-like measures had a slightly heavier upper tail (top 1% |*r_rm_*| = 0.118 against 0.066).

**Figure 3c** shows this as the trait-like cumulative curve trailing further to the right. Replication rates were similarly low in both sets, at 5.0% for the trait-like measures and 5.6% for the state-like measures (*r*_between_ = −0.010 and 0.015). As in the full analysis, no association survived FDR correction in either set (**Table S11**).

### 3.6 Interactive dashboard

To make these benchmarks directly usable, the full set of associations can be summarised through an interactive dashboard at https://ukbbmh-dashboard.up.railway.app/. Two views are provided, one for the cross-sectional Pearson’s correlations and one for the within-subject repeated measures correlations. In each, users select any combination of mental health domains or categories and any combination of imaging modalities or measure groups (**Table 2**), whether a single category against a single measure group, every category within a domain, or categories and measures drawn from different domains and modalities, and the results are summarised at any of these four levels. The longitudinal view can additionally restrict MHPs to trait-like or state-like measures. Each selection returns the four metrics used throughout this paper, the number of variables available, median |*r*|, top 1% |*r*|, and the replication rate, plotted across the groups being compared. Far too many selections exist to tabulate in advance, so these values are computed from the underlying associations for each request, extending the category- and modality-level summaries reported here (**Figure 2**; **Tables S7–9**) to any subset of phenotypes a study might consider. Everything plotted, together with a table of the underlying values and a record of the filters applied, can be downloaded as a self-contained HTML report. This allows the variable availability, expected effect size, and replicability of any selection of mental health and imaging phenotypes to be read off directly at the point of designing a study.

## 4 Discussion

In this study, we set out to establish benchmarks for brain-mental health associations in the UKB, estimating both the magnitude and the replicability of univariate correlations across 17,580 IDPs and 309 MHPs, resolved by mental health category, imaging modality, and, where repeated measurements allowed, examined longitudinally. The associations were small throughout and only modestly replicated, a pattern that held across every subset of associations we examined. Within that consistently small landscape, however, the effects were not uniform. Some mental health categories and imaging modalities yielded systematically larger or more replicable associations than others, and magnitude and replicability did not always track together, such that the largest effects were often among the least replicable.

### 4.1 A small overall effect-size landscape

Overall, the univariate brain-mental health associations are very small, and the median effect was an order of magnitude weaker than the effects routinely reported in smaller neuroimaging studies. Only associations at the extreme end of the distribution reached what would conventionally be called a small-to-moderate effect. The same pattern held when the analysis was repeated in a separate group of participants, indicating that this is a stable property of the data rather than a feature of any one participant subset.

These magnitudes are broadly consistent with existing UKB benchmarks. Dutt et al. (2022) reported median absolute correlations in the order of 0.01 to 0.04 for five summary scores of depression and anxiety. Our overall median sits toward the lower end of this range, as expected given our broader and more heterogeneous phenotype coverage. Miller et al. (2016), in their foundational characterisation of the imaging resource, correlated IDPs against roughly 1,100 non-imaging variables and reported maximum univariate effects approaching |*r*| = 0.25. Those non-imaging variables spanned lifestyle factors such as diet and exercise, physical and anthropometric measures such as body mass index and bone density, and cognitive test scores. The present analysis adds mental health to that landscape, and its top 1% associations reach into and beyond the upper bound set by the lifestyle, physical, and cognitive measures already characterised.

This pattern of very small univariate effects is what the BWAS framework describes. Marek et al. (2022) reported a median univariate effect of |*r*| = 0.01 and top 1% of |*r*| = 0.06 in the Adolescent Brain Cognitive Development (ABCD) cohort, and we reported similarly small effects in an older adult population. This establishes that these small effects generalise across the lifespan rather than being specific to developmental samples. Marek et al. (2022) showed that even their largest split-half analyses (*n* = 1,964) failed to deliver high replication, with only 25% of nominally significant associations being replicated. Our subsets are more than twice that size, yet the replication rate was even lower. Our findings therefore show that replication remains poor at sample sizes in the thousands, a result that aligns with the conclusion of Marek et al. (2022) that replicable brain-wide associations require very large samples and that even very large samples may be insufficient when effects are this small.

An association needed only to exceed |*r*| of about 0.07 to become statistically significant, which corresponds to a brain measure accounting for roughly half a percent of the variance in a mental health measure. This challenge of testing at population scale is acknowledged by Miller et al. (2016), who noted that even correlations as small as *r* ≈ 0.1, explaining about 1% of population variance, became significant despite correction for multiple comparisons. A statistically significant association in the UKB can therefore reflect an effect far too small to be of practical interest.

Replication fell steadily as effect size rose, so that the largest observed correlations replicated least often. This is the pattern commonly referred to as the winner’s curse (Button et al., 2013). When the true associations cluster near zero, the largest observed values are disproportionately those inflated by chance, and this inflation is worst for effects that only cross the significance threshold because they happen to be overestimated. The strongest observed correlations are therefore also the least likely to replicate in an independent subset. The same logic explains a result that might otherwise seem counterintuitive: associations surviving FDR correction replicated no more often than nominally significant ones, and in fact slightly less often (6.9% against 9.0%). This is consistent with Marek et al. (2022), who found that effect-size inflation among statistically significant associations is greater under stricter correction (*p* < 10⁻⁷) than under a lenient threshold (*p* < 0.05), so that tightening the threshold preferentially retains the very correlations most exposed to inflation. The strongest individual associations are thus the ones that warrant the most caution, and no single association should be regarded as reliable until it has been observed again in separate groups of participants.

### 4.2 Variation across mental health categories

Depression and anxiety dominate prior brain-mental health research in the UKB (Davis, Mirza, et al., 2025), and their univariate brain associations are small, in line with the modest correlations Dutt et al. (2022) reported for summary depression and anxiety scores. Because these are the categories researchers most often study, our benchmarks bear directly on how their findings should be read. Large individual associations reported in any study should be taken to belong to the upper tail of an otherwise small distribution, and may not replicate even in samples of several thousand, whereas the smaller and more consistent effects sometimes regarded as uninformative sit closer to the realistic population magnitude and should not be discounted.

Among imaging modalities, tfMRI produced the largest effects for BD and depression of any modality apart from ASL. This is unsurprising, since the tfMRI protocol uses an emotion face-matching task, which probes emotional processing directly. Tamm et al. (2022) provide a case where our benchmarks change how a single result should be read. In more than 28,000 UKB participants, they found that amygdala reactivity during this paradigm, the most established tfMRI correlate of depression, explained essentially none of the variance in concurrent depressive symptoms. The same phenotype measures enter our analysis, where their associations with depression sit in the middle of a distribution whose typical value is around |*r*| = 0.016. The near-null amygdala association is therefore consistent with the broader pattern observed across brain measures, where effects of this magnitude are typical rather than exceptional.

Resolving the analysis to regional IDPs shows where in the cortex an association lies, illustrated here by cortical thickness and the depression phenotypes. Cortical thickness was most consistently related to the depression phenotypes in the bilateral pars triangularis and pars opercularis, the left postcentral cortex, and the right superior parietal and lateral occipital cortex. Meta-analysis of clinical case-control studies has localized cortical thinning in major depression to the left pars opercularis among other regions (Suh et al., 2019), so the population-scale association we found overlaps with a pattern already established in clinical samples.

The most-studied transdiagnostic features in the UKB are self-harm thoughts and behaviours (Davis, Mirza, et al., 2025), which yielded the largest median and upper-tail effects within the domain, yet replicated least often across participant subsets. Closely tied to both depression and anxiety, self-harm produced effects approaching the magnitude of those observed for the mental disorder categories, indicating that it warrants inclusion as an additional variable of interest in studies of brain-mental health relationships. Neuroticism, the other widely studied feature in UKB neuroimaging research (Dutt et al., 2022), produced effects in line with previously reported estimates (on the order of |*r*| ≈ 0.01–0.03). Its replication rate, while comparatively higher within the transdiagnostic domain, remained modest in absolute terms, underscoring that even well-studied personality traits with consistent prior findings yield brain associations that are small and replicate inconsistently across samples.

ED emerged as the category with the largest effects in the analysis, and as the clearest priority for follow-up. Decomposing the category by modality locates much of the effect in dMRI, which after ASL was both the strongest and the most replicable modality here, implicating alterations in white matter microstructure. This accords with the clinical literature on anorexia nervosa, where a voxel-based meta-analysis of 13 diffusion-tensor imaging studies (227 patients, 243 controls) found reduced FA across several white matter tracts in patients relative to controls (Barona et al., 2019), indicating compromised white matter integrity. T1-based structural measures may also merit attention in ED, even though they did not produce the largest modality-specific effect in our analysis. Cortical and subcortical alterations in anorexia nervosa are among the largest observed in any psychiatric condition, exceeding those reported for depression, anxiety, and other common disorders (Walton et al., 2022). Because items of ED were added to the cohort only with the MWB 2022+ questionnaire (Davis, Coleman, et al., 2025), no prior UKB study has related them to brain measures.

### 4.3 Variation across imaging modalities

A similar picture emerged across imaging modalities. Effects were small and broadly similar across the modalities that dominate the UKB protocol. T1, dMRI, and rfMRI together account for the vast majority of IDPs and are the modalities most widely used in mental health research, since all three support dense, brain-wide measurement, capturing cortical and subcortical morphology at the regional level, white matter microstructure at the tract level, and intrinsic functional activity at the node and network level. Each produced median and top 1% effects close to the overall average, with replication rates of 8.8–9.4%. As with the mental health categories, large individual associations in these modalities again fall in the upper tail of an otherwise small distribution and warrant the same caution.

The remaining modalities each contribute far fewer IDPs, so their estimates rest on a smaller set of variables and should be read with that in mind. ASL produced the largest median and top 1% effects of any modality. It indexes cerebral perfusion, a physiologically distinct signal that structural and BOLD measures do not capture (Haller et al., 2016). These effects should be read as preliminary. ASL IDPs have been available in UKB only since 2023, so the ASL associations were estimated from a smaller subsample than the other modalities, and the reduced sample size plausibly inflates the absolute effect sizes while leaving replication moderate. The magnitude of these effects, together with the limited use of perfusion measures in UKB mental health research to date, nonetheless marks ASL as a priority target for hypothesis-driven follow-up. SWI measures magnetic susceptibility in deep grey-matter structures and is sensitive to tissue properties such as iron content (Wang et al., 2022). Its effects were comparable in size to the other modalities, but it replicated more consistently than any of them. At 18.4%, however, this replication rate remained modest in absolute terms. T2 contributes the smallest IDP set of any modality, yet 11.9% of its associations were nominally significant, nearly double the 6.3% across all IDPs, and 14.6% replicated. Although these proportions are computed over 1,854 correlations and are less stable than those for the dominant modalities, white matter hyperintensity burden has been linked to depression and anxiety in earlier work (Wang et al., 2014; Zhou et al., 2024). Across mental health categories examined here, it replicated most often for subjective well-being, which merits follow-up.

### 4.4 Longitudinal stability

The same-day associations showed different patterns at the two imaging visits. The initial visit produced smaller effects that replicated more often, whereas the repeat visit produced larger but less replicable effects. Taken on its own, this difference could be read as the associations strengthening over the roughly five years separating the visits, but the cross-visit associations show otherwise. Pairing IDPs from one visit with MHPs from the other barely shifted the distribution, so what set the effect sizes and replication rates was which visit supplied the IDPs, not the interval between the brain and mental health measurements, and not which of the two was assessed first. Separations of this length, five years at the median and ranging from one to ten years, therefore left both the magnitude and the replicability of the associations nearly unchanged. Cross-sectional studies in UKB frequently relate brain and mental health measures collected at different assessment waves, and our results suggest this practice comes at little cost to the size of the associations detected. The difference that does remain between the two visits likely reflects the properties of the imaging data rather than the passage of time. The repeat imaging visit has more missing data than the initial visit, which reduces the number of participants contributing to each correlation, and these smaller samples inflate absolute effect sizes while reducing their replicability across the discovery and validation subsets.

Repeated-measures correlations remove between-subject variance to ask whether brain and mental health change together within the same individual across the two imaging visits. These correlations centred near zero and none survived correction. Age was deliberately retained rather than partialled out, so any coordinated age-related change in brain and mental health would have inflated these correlations. That they remained near zero therefore indicates that the null result is not an artefact of over-adjustment. The most straightforward interpretation is that, over intervals of around five years in this older adult sample, change within individuals in both brain and mental health measures is too small relative to measurement error for any association to emerge. This limit is well documented for brain measures, where standard longitudinal MRI estimates can fail to track within-subject change because the measurement error is often as large as the change itself (Elliott et al., 2026).

Splitting the mental health measures by how much they can plausibly change within a person did not change this picture. The trait-like measures, comprising lifetime symptoms, diagnoses, and personality traits, showed a marginally heavier upper tail than the state-like measures of recent symptoms, mood, and circumstances, but this difference is slight and both distributions sit close to zero. Replication stayed flat across the full range of effect sizes rather than rising with them, so the heavier trait-like tail reflects the noisier estimates expected when within-subject variance is small. For most questions about brain-mental health relationships in UKB, between-subject analyses are therefore the more informative use of the dataset at present, with within-subject correlations reserved for measures where meaningful change is plausible. But as UKB acquires further imaging data, the longer intervals should raise the ratio of genuine change to measurement error, so that within-subject analyses may become more viable than they are at present.

### 4.5 Limitations

Several limitations should be considered. The UKB cohort is not representative of the UK or global population, and the sub-cohort contributing to our analyses amplifies this bias, since it is restricted to participants who returned for a second imaging visit and completed both online questionnaires. MHQ respondents are already less likely to live in deprived areas or report a longstanding illness than the broader cohort (Davis, Coleman, et al., 2025), and individuals with severe mental illness are under-represented (Davis, Mirza, et al., 2025). The benchmarks reported here therefore describe associations in a relatively healthy, mostly Caucasian, older adult sample, and their magnitude and replicability may differ in clinical or more diverse populations. We also restricted the analysis to self-report questionnaire data and set aside the linked National Health Service diagnostic records and polygenic scores available in UKB. Self-report items capture current and lifetime symptoms, so associations built on ascertained diagnoses may behave differently, and extending these benchmarks to the record-linkage phenotypes is a natural complement to the present work. Also, the test-retest reliability of the phenotypes places a ceiling on the correlations that can be detected, and this ceiling varies across measures. Test-retest agreement in UKB ranges from substantial for some items to modest for others, for example from κ = 0.81 for lifetime cannabis use to κ = 0.30 for lifetime BD (Davis, Coleman, et al., 2025), so the weaker associations we report for less consistent phenotypes partly reflect limited reliability rather than a genuine absence of signal. Finally, our univariate analyses do not capture multivariate brain-mental health relationships, which Dutt et al. (2022) and Marek et al. (2022) have shown can be larger in size and more replicability. The benchmarks reported here should therefore be read as a lower bound on what can be detected in the UKB, not as a ceiling on the brain-mental health relationships the dataset can support.

### 4.6 Future implications

The benchmarks established here are intended as a practical resource for planning future brain-mental health research. For researchers intending to work with the UKB, they support study scoping and variable selection. In choosing which mental health categories and imaging modalities to pursue, magnitude and replicability should be weighed together rather than separately. The categories and modalities with the largest effects were often not those that replicated most reliably, and selecting on either property alone risks chasing associations that will not hold or discarding smaller effects that sit closer to the true population magnitude. For those planning studies outside the UKB, whether for large-scale cohorts or smaller clinical samples, the estimates reported here provide population-scale effect sizes to anchor power calculations. Because they are drawn from a population cohort, they avoid the inflation carried by estimates from smaller or clinically enriched samples, which overstate what is detectable and leave studies underpowered for the effects realistically present. Both uses are served by an interactive dashboard at https://ukbbmh-dashboard.up.railway.app/, which returns these benchmarks for any selection of mental health categories and imaging modalities. Because magnitude and replicability are returned side by side for the same selection, the trade-off between them can be inspected directly for whichever categories and modalities a study is considering.

The following priorities follow from these benchmarks, identifying where new studies in the UKB are most urgently needed. ED showed the largest associations of any category yet have never been related to brain measures in the UKB, making them the clearest priority. The dMRI IDPs are the natural starting point, as diffusion was both the strongest and the most replicable modality for this category, with the T1 measures a reasonable complement given the structural alterations documented in anorexia nervosa. For depression and BD, tfMRI produced the largest effects after ASL. Work with these IDPs has centred on amygdala reactivity, where the absence of an association with depressive symptoms reflects an effect size typical of what we report throughout rather than an unusually weak one, so the remaining tfMRI IDPs are an open target for studies of mood disorders. Two modalities are underused relative to what they yielded here. ASL yielded the largest effects of any modality but is little used in UKB mental health research, marking it as a candidate for mood disorders and self-harm, though estimates drawn from a smaller subsample are better suited to generating hypotheses. SWI replicated more consistently than any other modality, most notably for substance use, and can be prioritised where replicability of an association matters more than its size. Whichever category-modality pairing is chosen, the regional IDPs allow it to be localised across the cortex, subcortical structures, and white matter tracts, as shown here for cortical thickness and depression. Longitudinal work should first confirm that the chosen phenotypes vary meaningfully within individuals over the interval available, since within-subject change over a median of five years proved too small to yield detectable associations.

In summary, brain-mental health associations in the UKB are small and replicate poorly. The categories and modalities with the largest effects were often not those that held up in an independent subset, so neither magnitude nor replicability can stand in for the other when choosing what to study. Separating the brain and mental health measurements in time did not weaken their association, and within-subject change over a median of five years was too small to yield detectable associations. Power calculations anchored to these population-scale estimates give a study a fair chance of detecting what is actually there. The interactive dashboard makes the estimates available for any category-modality combination a study might consider.

## Data Availability

This research was conducted using the UK Biobank Resource under application number 100773. UK Biobank data are available to bona fide researchers through an application to UK Biobank (https://www.ukbiobank.ac.uk/enable-your-research). The summary statistics reported here, comprising effect size, significance, and replication measures for all associations, are provided in Supplementary Tables S7–S11. Summaries over any selection of mental health categories and imaging modalities can be generated interactively at https://ukbbmh-dashboard.up.railway.app/.

## Code Availability

Code for the analyses reported here is available at https://github.com/XuqianLi5225/ukbbmh-mhp.git.

## Ethics

UK Biobank holds Research Tissue Bank approval from the North West Multi-centre Research Ethics Committee (REC reference 21/NW/0157), under which researchers may use the resource without further ethical approval. All participants provided written informed consent at recruitment. This research was conducted under UK Biobank application number 100773.

## Acknowledgements

We thank the UK Biobank participants and staff. This work was supported by an NHMRC Investigator grant (grant number: 2007718) and AIBN start-up funding to L.O. S.K. was supported by a Postgraduate Research Scholarship from the University of Queensland.

## Author Contributions

X.L., S.K., and L.O. conceived the study. X.L. curated the data, developed the analysis code and the web application, performed the analyses, produced the figures, administered the project, and wrote the original draft. S.K. contributed to data curation and validated the analyses. L.O. contributed to data curation, provided resources, validated the analyses, and supervised the project. All authors contributed to the methodology and reviewed and edited the manuscript.

## Notes

https://ukbbmh-dashboard.up.railway.app/

### Competing Interest Statement

The authors have declared no competing interest.

