## Supplementary material for "An updated roadmap for brain-mental health associations in the UK Biobank": Table S1

### Supplementary Materials

**Table S1. Official UKB Resources, Documentation, and Publications for Mental Health Research**

| Questionnaire | Wave | Type | Reference | Description | URL |
|---|---|---|---|---|---|
| T/Screen 2009+ | 0–3 | Documentation | Resource 100247 | Procedure for administering the T/Screen 2009+ | https://biobank.ndph.ox.ac.uk/showcase/ukb/docs/Touchscreen.pdf |
| | 0–3 | Documentation | Resource 113241 | Content of the enhanced T/Screen 2009+ | https://biobank.ndph.ox.ac.uk/showcase/ukb/docs/TouchscreenQuestionsMainFinal.pdf |
| | 0 | Webpage | / | Timeline of baseline data collection | https://biobank.ndph.ox.ac.uk/ukb/exinfo.cgi?src=baseline_data |
| | 0 | Publication | Smith et al., 2013 | Methodologies and results from T/Screen 2009+ in 172,751 participants | https://doi.org/10.1371/journal.pone.0075362 |
| | 0–3 | Documentation | Resource 158772 | List of derived data-fields describing the status of major depression, bipolar disorder, and neuroticism based on methodologies in Smith et al. (2013) | https://biobank.ndph.ox.ac.uk/showcase/ukb/docs/MentalStatesDerivation.pdf |
| | 1 | Webpage | / | Overview of the repeat assessment visit | https://biobank.ndph.ox.ac.uk/ukb/exinfo.cgi?src=repeat_assessment_2012 |
| | 2–3 | Webpage | / | Overview of both imaging visits | https://www.ukbiobank.ac.uk/enable-your-research/about-our-data/imaging-data |
| | 2 | Publication | Littlejohns et al., 2020 | Rationales and methodologies of the UKB imaging study | https://doi.org/10.1038/s41467-020-15948-9 |
| | 3 | Webpage | / | Overview of the repeat imaging visit | https://www.ukbiobank.ac.uk/learn-more-about-uk-biobank/news/world-s-largest-imaging-study-begins-exciting-second-stage-to-re-scan-60-000-uk-biobank-volunteers |
| | 3 | Documentation | Resource 120 | Selection criteria and timelines of the repeat imaging visit | https://biobank.ndph.ox.ac.uk/showcase/ukb/docs/first_repeat_imaging_selection.pdf |
| MHQ 2016+ | / | Documentation | Resource 22 | Overview and content of the online Mental Health Questionnaire | https://biobank.ndph.ox.ac.uk/showcase/ukb/docs/mental_health_online.pdf |
| | / | Publication | Davis & Hotopf, 2019 | Rationale and methodologies of mental health phenotyping in UKB | https://wileymicrositebuilder.com/progress/wp-content/uploads/sites/28/2019/02/Comment-Davis-final-lsw.pdf |

| Questionnaire | Wave | Type | Reference | Description | URL |
|---|---|---|---|---|---|
| | / | Publication | Davis et al., 2019 | Methodologies on classification of mental disorder from MHQ 2016+ in 157,363 participants | https://doi.org/10.1002/mpr.1796 |
| | / | Publication | Davis et al., 2020 | Methodologies and results from MHQ 2016+ in 157,366 participants | https://doi.org/10.1192/bjo.2019.100 |
| MWB 2022+ | / | Documentation | Resource 2800 | Overview and content of the online Mental Well-being Questionnaire | https://biobank.ndph.ox.ac.uk/showcase/ukb/docs/mwb_overview.pdf |
| | / | Publication | Davis, Coleman, et al., 2025 | The UK Biobank mental health enhancement 2022: Methods and results | https://doi.org/10.1371/journal.pone.0324189 |
| Multiple | / | Publication | Davis, Mirza, et al., 2025 | Unlocking mental health insights with UK Biobank data: Past use and future opportunities | https://doi.org/10.1017/S0033291725101359 |

*Note.* T/Screen = Touchscreen Questionnaire; MHQ = Mental Health Questionnaire; MWB = Mental Well-being Questionnaire.

**Table S2. Mental Health Variables Requiring Additional Missing Value Definitions**

| Field ID | Description | Assessment Waves | Value Type | Data Coding ID | Data Coding for Special Value |
|---|---|---|---|---|---|
| 20420 | Longest period spent worried or anxious | MHQ 2016+ | Integer | 517 | -999: All my life / as long as I can remember |
| 20442 | Lifetime number of depressed periods | MHQ 2016+ | Integer | 511 | -999: Too many to count / One episode ran into the next |
| 20461 | Age when first had unusual or psychotic experience | MHQ 2016+ | Integer | 530 | -999: As long as I can remember |
| 20465 | Number of times heard an un-real voice | MHQ 2016+ | Integer | 528 | -999: Too many to count |
| 20470 | Number of times believed in an un-real conspiracy against self | MHQ 2016+ | Integer | 528 | -999: Too many to count |
| 20473 | Number of times seen an un-real vision | MHQ 2016+ | Integer | 528 | -999: Too many to count |
| 20476 | Number of times believed in un-real communications or signs | MHQ 2016+ | Integer | 528 | -999: Too many to count |
| 29033 | Lifetime number of depressed periods | MWB 2022+ | Integer | 1907 | -4: Too many to count / One episode ran into the next |
| 29055 | Lifetime number of manic or irritable periods | MWB 2022+ | Integer | 1924 | -4: Too many to count / One episode ran into the next |
| 29071 | Lifetime number of periods of worry lasting one month or more | MWB 2022+ | Integer | 1912 | -4: Too many to remember |
| 29125 | Lowest weight during period when underweight | MWB 2022+ | Integer, kg | 1955 | 0: Invalid weight |
| 29170 | When retired | MWB 2022+ | Integer, calendar year | 1929 | 0: I am still working in some capacity;<br>1: Before 2017 |

*Note.* T/Screen = Touchscreen Questionnaire; MHQ = Mental Health Questionnaire; MWB = Mental Well-being Questionnaire.

**Table S3. Mental Health Variables with Data Codes Being Reversed**

| Field ID | Description | Assessment Waves | Data Coding ID | Data Coding | Reversed Data Coding |
|---|---|---|---|---|---|
| 1031 | Frequency of friend/family visits | T/Screen 2009+ 0–3 | 100327 | 1: Almost daily;<br>2: 2-4 times a week;<br>3: About once a week;<br>4: About once a month;<br>5: Once every few months;<br>6: Never or almost never;<br>7: No friends/family outside household;<br>-1: Do not know;<br>-3: Prefer not to answer | 7: Almost daily;<br>6: 2-4 times a week;<br>5: About once a week;<br>4: About once a month;<br>3: Once every few months;<br>2: Never or almost never;<br>1: No friends/family outside household |
| 4526 | Happiness | T/Screen 2009+ 0–3 | 100478 | 1: Extremely happy;<br>2: Very happy;<br>3: Moderately happy;<br>4: Moderately unhappy;<br>5: Very unhappy;<br>6: Extremely unhappy;<br>-1: Do not know;<br>-3: Prefer not to answer | 6: Extremely happy;<br>5: Very happy;<br>4: Moderately happy;<br>3: Moderately unhappy;<br>2: Very unhappy;<br>1: Extremely unhappy |
| 4537 | Work/job satisfaction | T/Screen 2009+ 0–3 | 100479 | 1: Extremely happy;<br>2: Very happy;<br>3: Moderately happy;<br>4: Moderately unhappy;<br>5: Very unhappy;<br>6: Extremely unhappy;<br>7: I am not employed;<br>-1: Do not know;<br>-3: Prefer not to answer | 7: Extremely happy;<br>6: Very happy;<br>5: Moderately happy;<br>4: Moderately unhappy;<br>3: Very unhappy;<br>2: Extremely unhappy;<br>1: I am not employed |
| 4548 | Health satisfaction | T/Screen 2009+ 0–3 | 100478 | 1: Extremely happy;<br>2: Very happy;<br>3: Moderately happy;<br>4: Moderately unhappy;<br>5: Very unhappy; | 6: Extremely happy;<br>5: Very happy;<br>4: Moderately happy;<br>3: Moderately unhappy;<br>2: Very unhappy; |

| Field ID | Description | Assessment Waves | Data Coding ID | Data Coding | Reversed Data Coding |
|---|---|---|---|---|---|
| | | | | 6: Extremely unhappy;<br>-1: Do not know;<br>-3: Prefer not to answer | 1: Extremely unhappy |
| 4559 | Family relationship satisfaction | T/Screen 2009+<br>0–3 | 100478 | 1: Extremely happy;<br>2: Very happy;<br>3: Moderately happy;<br>4: Moderately unhappy;<br>5: Very unhappy;<br>6: Extremely unhappy;<br>-1: Do not know;<br>-3: Prefer not to answer | 6: Extremely happy;<br>5: Very happy;<br>4: Moderately happy;<br>3: Moderately unhappy;<br>2: Very unhappy;<br>1: Extremely unhappy |
| 4570 | Friendships satisfaction | T/Screen 2009+<br>0–3 | 100478 | 1: Extremely happy;<br>2: Very happy;<br>3: Moderately happy;<br>4: Moderately unhappy;<br>5: Very unhappy;<br>6: Extremely unhappy;<br>-1: Do not know;<br>-3: Prefer not to answer | 6: Extremely happy;<br>5: Very happy;<br>4: Moderately happy;<br>3: Moderately unhappy;<br>2: Very unhappy;<br>1: Extremely unhappy |
| 4581 | Financial situation satisfaction | T/Screen 2009+<br>0–3 | 100478 | 1: Extremely happy;<br>2: Very happy;<br>3: Moderately happy;<br>4: Moderately unhappy;<br>5: Very unhappy;<br>6: Extremely unhappy;<br>-1: Do not know;<br>-3: Prefer not to answer | 6: Extremely happy;<br>5: Very happy;<br>4: Moderately happy;<br>3: Moderately unhappy;<br>2: Very unhappy;<br>1: Extremely unhappy |
| 10740 | Frequency of friend/family visits (pilot) | T/Screen 2009+<br>Pilot | 100662 | 1: Almost daily;<br>2: 2-4 times a week;<br>3: About once a week;<br>4: About once a month;<br>5: Once every few months;<br>6: Never or almost never; | 7: Almost daily;<br>6: 2-4 times a week;<br>5: About once a week;<br>4: About once a month;<br>3: Once every few months;<br>2: Never or almost never; |

| Field ID | Description | Assessment Waves | Data Coding ID | Data Coding | Reversed Data Coding |
|---|---|---|---|---|---|
| | | | | 7: No friends/family outside household;<br>-3: Prefer not to answer | 1: No friends/family outside household |
| 20458 | General happiness | MHQ 2016+ | 537 | -818: Prefer not to answer;<br>-121: Do not know;<br>1: Extremely happy;<br>2: Very happy;<br>3: Moderately happy;<br>4: Moderately unhappy;<br>5: Very unhappy;<br>6: Extremely unhappy | 6: Extremely happy;<br>5: Very happy;<br>4: Moderately happy;<br>3: Moderately unhappy;<br>2: Very unhappy;<br>1: Extremely unhappy |
| 20459 | General happiness with own health | MHQ 2016+ | 537 | -818: Prefer not to answer;<br>-121: Do not know;<br>1: Extremely happy;<br>2: Very happy;<br>3: Moderately happy;<br>4: Moderately unhappy;<br>5: Very unhappy;<br>6: Extremely unhappy | 6: Extremely happy;<br>5: Very happy;<br>4: Moderately happy;<br>3: Moderately unhappy;<br>2: Very unhappy;<br>1: Extremely unhappy |
| 29014 | Fraction of day affected during worst episode of depression | MWB 2022+ | 1935 | 0: All day long;<br>1: Most of the day;<br>2: About half of the day;<br>3: Less than half of the day;<br>-1: Do not know;<br>-3: Prefer not to answer | 3: All day long;<br>2: Most of the day;<br>1: About half of the day;<br>0: Less than half of the day |
| 29015 | Frequency of depressed days during worst episode of depression | MWB 2022+ | 1936 | 0: Every day;<br>1: Almost every day;<br>2: Less often;<br>-1: Do not know;<br>-3: Prefer not to answer | 2: Every day;<br>1: Almost every day;<br>0: Less often |

| Field ID | Description | Assessment Waves | Data Coding ID | Data Coding | Reversed Data Coding |
|---|---|---|---|---|---|
| 29031 | Impact on normal roles during worst period of depression | MWB 2022+ | 1946 | 0: A lot;<br>1: Somewhat;<br>2: A little;<br>3: Not at all;<br>-3: Prefer not to answer | 3: A lot;<br>2: Somewhat;<br>1: A little;<br>0: Not at all |
| 29133 | Longest amount of time that overate or binge ate at least once a week | MWB 2022+ | 1919 | 0: At least 3 months;<br>1: More than 1 month but less than 3 months;<br>2: Less than 1 month;<br>-3: Prefer not to answer | 2: At least 3 months;<br>1: More than 1 month but less than 3 months;<br>0: Less than 1 month |
| 29135 | Frequency of feeling had no control overeating during episodes of overeating/binge eating | MWB 2022+ | 1921 | 0: At least once a week for at least 3 months;<br>1: At least once a week for between 1 and 3 months;<br>2: Occasionally;<br>3: Never;<br>-3: Prefer not to answer | 3: At least once a week for at least 3 months;<br>2: At least once a week for between 1 and 3 months;<br>1: Occasionally;<br>0: Never |
| 29142 | Longest amount of time that engaged in these behaviours when overeating/binge eating | MWB 2022+ | 1919 | 0: At least 3 months;<br>1: More than 1 month but less than 3 months;<br>2: Less than 1 month;<br>-3: Prefer not to answer | 2: At least 3 months;<br>1: More than 1 month but less than 3 months;<br>0: Less than 1 month |
| 29145 | Longest amount of time that engaged in these behaviours outside of periods of low weight or overeating/binge eating | MWB 2022+ | 1919 | 0: At least 3 months;<br>1: More than 1 month but less than 3 months;<br>2: Less than 1 month;<br>-3: Prefer not to answer | 2: At least 3 months;<br>1: More than 1 month but less than 3 months;<br>0: Less than 1 month |
| 29161 | Recovery from COVID-19 | MWB 2022+ | 1902 | 0: Yes, completely;<br>1: Yes, mostly;<br>2: Partially;<br>3: No, not at all;<br>4: No, getting worse; | 4: Yes, completely;<br>3: Yes, mostly;<br>2: Partially;<br>1: No, not at all;<br>0: No, getting worse |

| Field ID | Description | Assessment Waves | Data Coding ID | Data Coding | Reversed Data Coding |
|---|---|---|---|---|---|
| | | | | -1: Do not know;<br>-3: Prefer not to answer | |
| 29181 | General happiness | MWB 2022+ | 100478 | 1: Extremely happy;<br>2: Very happy;<br>3: Moderately happy;<br>4: Moderately unhappy;<br>5: Very unhappy;<br>6: Extremely unhappy;<br>-1: Do not know;<br>-3: Prefer not to answer | 6: Extremely happy;<br>5: Very happy;<br>4: Moderately happy;<br>3: Moderately unhappy;<br>2: Very unhappy;<br>1: Extremely unhappy |

*Note.* T/Screen = Touchscreen Questionnaire; MHQ = Mental Health Questionnaire; MWB = Mental Well-being Questionnaire.

**Table S4. Mental Health Variables with Multiple Selections**

| Field ID | Description | Assessment Waves | Data Coding ID | Data Coding | Range of New Coding |
|---|---|---|---|---|---|
| 6145 [a] | Illness, injury, bereavement, stress in last 2 years | T/Screen 2009+ 0–3 | 100502 | 1: Serious illness, injury or assault to yourself;<br>2: Serious illness, injury or assault of a close relative;<br>3: Death of a close relative;<br>4: Death of a spouse or partner;<br>5: Marital separation/divorce;<br>6: Financial difficulties;<br>-7: None of the above;<br>-3: Prefer not to answer | 0–6 |
| 6156 [a, b] | Manic/hyper symptoms | T/Screen 2009+ 0–3 | 100498 | 11: I was more active than usual;<br>12: I was more talkative than usual;<br>13: I needed less sleep than usual;<br>14: I was more creative or had more ideas than usual;<br>15: All of the above;<br>-7: None of the above | 0–4 |
| 6160 [a] | Leisure/social activities | T/Screen 2009+ 0–3 | 100328 | 1: Sports club or gym;<br>2: Pub or social club;<br>3: Religious group;<br>4: Adult education class;<br>5: Other group activity;<br>-7: None of the above;<br>-3: Prefer not to answer | 0–5 |
| 10721 [a] | Illness, injury, bereavement, stress in last 2 years (pilot) | T/Screen 2009+ Pilot | 100683 | 1: Serious illness, injury or assault to yourself;<br>2: Serious illness, injury or assault of a close relative;<br>3: Death of a close relative;<br>4: Death of a spouse or partner;<br>5: Marital separation/divorce;<br>6: Financial difficulties;<br>-7: None of the above;<br>-1: Do not know;<br>-3: Prefer not to answer | 0–6 |

| Field ID | Description | Assessment Waves | Data Coding ID | Data Coding | Range of New Coding |
|---|---|---|---|---|---|
| 20544 | Mental health problems ever diagnosed by a professional | MHQ 2016+ | 1401 | 1: Social anxiety or social phobia;<br>2: Schizophrenia;<br>3: Any other type of psychosis or psychotic illness;<br>4: A personality disorder;<br>5: Any other phobia (e.g., disabling fear of heights or spiders);<br>6: Panic attacks;<br>7: Obsessive compulsive disorder (OCD);<br>10: Mania, hypomania, bipolar or manic-depression;<br>11: Depression;<br>12: Bulimia nervosa;<br>13: Psychological over-eating or binge-eating;<br>14: Autism, Asperger's or autistic spectrum disorder;<br>15: Anxiety, nerves or generalized anxiety disorder;<br>16: Anorexia nervosa;<br>17: Agoraphobia;<br>18: Attention deficit or attention deficit and hyperactivity disorder (ADD/ADHD);<br>-818: Prefer not to answer (group A);<br>-819: Prefer not to answer (group B) | 1–16 |
| 20546 | Substances taken for depression | MHQ 2016+ | 1405 | 1: Unprescribed medication (more than once);<br>3: Medication prescribed to you (for at least two weeks);<br>4: Drugs or alcohol (more than once);<br>-818: Prefer not to answer | 1–3 |
| 20547 | Activities undertaken to treat depression | MHQ 2016+ | 1406 | 1: Talking therapies, such as psychotherapy, counselling, group therapy or CBT;<br>3: Other therapeutic activities such as mindfulness, yoga or art classes;<br>-818: Prefer not to answer | 1–2 |
| 20548 | Manifestations of mania or irritability | MHQ 2016+ | 1407 | 1: I was more talkative than usual;<br>2: I was more restless than usual;<br>3: My thoughts were racing;<br>5: I needed less sleep than usual;<br>6: I was more creative or had more ideas than usual; | 1–8 |

| Field ID | Description | Assessment Waves | Data Coding ID | Data Coding | Range of New Coding |
|---|---|---|---|---|---|
| | | | | 7: I was easily distracted;<br>8: I was more confident than usual;<br>9: I was more active than usual;<br>-818: Prefer not to answer | |
| 20549 | Substances taken for anxiety | MHQ 2016+ | 1405 | 1: Unprescribed medication (more than once);<br>3: Medication prescribed to you (for at least two weeks);<br>4: Drugs or alcohol (more than once);<br>-818: Prefer not to answer | 1–3 |
| 20550 | Activities undertaken to treat anxiety | MHQ 2016+ | 1406 | 1: Talking therapies, such as psychotherapy, counselling, group therapy or CBT;<br>3: Other therapeutic activities such as mindfulness, yoga or art classes;<br>-818: Prefer not to answer | 1–2 |
| 20551 | Substance of prescription or over-the-counter medication addiction | MHQ 2016+ | 1414 | 1: A sedative, benzodiazepine or sleeping tablet;<br>2: A painkiller;<br>3: Something else;<br>4: Do not know;<br>-818: Prefer not to answer | 1–3 |
| 20552 | Behavioural and miscellaneous addictions | MHQ 2016+ | 1415 | 1: Something else not mentioned;<br>2: A behaviour;<br>-818: Prefer not to answer | 1–2 |
| 20553 | Methods of self-harm used | MHQ 2016+ | 1422 | 1: Something not listed;<br>2: Swallowing dangerous objects or products;<br>3: Stopping prescribed medication;<br>4: Ingesting a medication in excess of the normal dose;<br>5: Self-injury such as self-cutting, scratching or hitting, etc.;<br>6: Ingesting alcohol or a recreational or illicit drug;<br>-818: Prefer not to answer | 1–6 |
| 20554 | Actions taken following self-harm | MHQ 2016+ | 1423 | 1: See anyone from psychiatric or mental health services, including liaison services;<br>3: Need hospital treatment (e.g., A&E);<br>4: Use a helpline/voluntary organization; | 1–5 |

| Field ID | Description | Assessment Waves | Data Coding ID | Data Coding | Range of New Coding |
| --- | --- | --- | --- | --- | --- |
| | | | | 5: See own GP;<br>6: Receive help from friends/family/neighbours;<br>-818: Prefer not to answer | |
| 29000 [a] | Mental health conditions ever diagnosed by a professional | MWB 2022+ | 1954 | -8: None of group C;<br>-7: None of group B;<br>-6: None of group A;<br>-5: Prefer not to answer group C;<br>-4: Prefer not to answer group B;<br>-3: Prefer not to answer group A;<br>1: Depression;<br>2: Mania, hypomania, bipolar or manic-depression;<br>3: Schizophrenia;<br>4: Any other type of psychosis or psychotic illness;<br>5: A personality disorder;<br>6: Autism, Asperger's or autistic spectrum disorder;<br>7: Attention deficit or attention deficit and hyperactivity disorder;<br>8: Obsessive compulsive disorder (OCD);<br>9: Anxiety or nerves;<br>10: Generalized anxiety disorder;<br>11: Social anxiety or social phobia<br>12: Agoraphobia;<br>13: Any other phobia (e.g. disabling fear of heights or spiders);<br>14: Panic attacks;<br>15: Panic disorder;<br>16: Post traumatic stress disorder (PTSD);<br>17: Anorexia nervosa;<br>18: Bulimia nervosa;<br>19: Binge-eating disorder;<br>20: Any other eating disorder | 0–20 |
| 29001 | Mental health conditions experienced by first degree blood relatives | MWB 2022+ | 1914 | 1: Depression;<br>2: Mania, hypomania, bipolar or manic-depression;<br>3: Schizophrenia; | 0–9 |

| Field ID | Description | Assessment Waves | Data Coding ID | Data Coding | Range of New Coding |
|---|---|---|---|---|---|
| | | | | 4: Any other type of psychosis or psychotic illness;<br>5: A personality disorder;<br>6: Autism, Asperger’s or autistic spectrum disorder;<br>7: Attention deficit or attention deficit and hyperactivity disorder;<br>8: An anxiety disorder;<br>9: An eating disorder;<br>0: None of the above;<br>-3: Prefer not to answer | |
| 29038 | Substances taken for depression | MWB 2022+ | 1956 | -3: Prefer not to answer;<br>0: None of options listed;<br>1: Drugs or alcohol (more than once);<br>2: Medication prescribed to participant (for at least 2 weeks);<br>3: Unprescribed medication (more than once) | 0–3 |
| 29039 | Use of specific medications | MWB 2022+ | 1908 | 1: Citalopram (sometimes called Cipramil);<br>2: Fluoxetine (Prozac or Oxactin);<br>3: Sertraline (Lustral);<br>4: Paroxetine (Seroxat);<br>5: Amitriptyline (Elavil);<br>6: Dosulepin (Prothiaden);<br>7: Other antidepressant(s);<br>-1: Do not know;<br>-3: Prefer not to answer | 1–7 |
| 29047 | Activities undertaken to treat depression | MWB 2022+ | 1957 | -3: Prefer not to answer;<br>0: None of options listed;<br>1: Other therapeutic activities such as mindfulness, yoga or art classes;<br>2: Talking therapies, such as psychotherapy, counselling, group therapy or CBT | 0–2 |
| 29051 | Manifestations of mania or irritability | MWB 2022+ | 1958 | -3: Prefer not to answer;<br>0: None of options listed;<br>1: More active than usual; | 0–8 |

| Field ID | Description | Assessment Waves | Data Coding ID | Data Coding | Range of New Coding |
|---|---|---|---|---|---|
| | | | | 2: More confident than usual;<br>3: Easily distracted;<br>4: More creative or had more ideas than usual;<br>5: Needed less sleep than usual;<br>6: Thoughts were racing;<br>7: More restless than usual;<br>8: More talkative than usual | |
| 29065 | Symptoms of panic attack | MWB 2022+ | 1910 | -3: Prefer not to answer;<br>0: No, I have not had this happen to me;<br>1: Heart was pounding or racing;<br>2: Sweating;<br>3: Trembling or shaking;<br>4: Felt short of breath, or like being smothered;<br>5: Felt like I was choking;<br>6: Pain or discomfort in chest;<br>7: Nauseous or felt sick in the stomach;<br>8: Felt dizzy, unsteady, light-headed or faint;<br>9: Felt hot or cold;<br>10: Felt numbness or tingling sensations;<br>11: Felt like things weren't real or detached from self;<br>12: Afraid of losing control or 'going crazy';<br>13: Afraid I was going to die | 0–13 |
| 29115 | Methods of self-harm used | MWB 2022+ | 1959 | -3: Prefer not to answer;<br>1: Ingesting alcohol or a recreational or illicit drug;<br>2: Self-injury such as self-cutting, scratching or hitting, etc.;<br>3: Ingesting a medication in excess of the normal dose;<br>4: Stopping prescribed medication;<br>5: Swallowing dangerous objects or products;<br>6: Something not listed | 1–6 |
| 29130 | Methods of controlling body shape or weight when at this low weight | MWB 2022+ | 1916 | 1: Made yourself vomit;<br>2: Used laxatives (pills or liquids);<br>3: Used diuretics (water pills);<br>4: Used weight loss pills; | 0–7 |

| Field ID | Description | Assessment Waves | Data Coding ID | Data Coding | Range of New Coding |
|---|---|---|---|---|---|
| | | | | 5: Exercised excessively or felt distressed if unable to exercise;<br>6: Fasted or not eaten for eight waking hours or more;<br>7: Other methods to lose weight/stay at low weight;<br>0: None of above;<br>-3: Prefer not to answer | |
| 29136 | Actions and feelings during periods of overeating/binge eating | MWB 2022+ | 1922 | 1: Eaten much more rapidly than normal;<br>2: Eaten until feeling uncomfortably full;<br>3: Eaten large amounts of food when not physically hungry;<br>4: Eaten alone because of feeling embarrassed by overeating;<br>5: Felt disgusted, depressed or very guilty afterward;<br>0: Done none of the above;<br>-3: Prefer not to answer | 0–5 |
| 29140 | Methods of controlling body shape or weight when overeating/binge eating | MWB 2022+ | 1916 | 1: Made yourself vomit;<br>2: Used laxatives (pills or liquids);<br>3: Used diuretics (water pills);<br>4: Used weight loss pills;<br>5: Exercised excessively or felt distressed if unable to exercise;<br>6: Fasted or not eaten for eight waking hours or more;<br>7: Other methods to lose weight/stay at low weight;<br>0: None of above;<br>-3: Prefer not to answer | 0–7 |
| 29167 | Sports and social activities attended in person at least once a week | MWB 2022+ | 1953 | 1: Sports club or gym/fitness class;<br>2: Pub or social club;<br>3: Religious group;<br>4: Adult education class;<br>5: Other group activity;<br>0: None of the above;<br>-3: Prefer not to answer | 0–5 |

| Field ID | Description | Assessment Waves | Data Coding ID | Data Coding | Range of New Coding |
|---|---|---|---|---|---|
| 29168 | Sports and social activities attended virtually at least once a week | MWB 2022+ | 1953 | 1: Sports club or gym/fitness class;<br>2: Pub or social club;<br>3: Religious group;<br>4: Adult education class;<br>5: Other group activity;<br>0: None of the above;<br>-3: Prefer not to answer | 0–5 |
| 29169 | Current situation | MWB 2022+ | 1928 | 1: Employed;<br>2: Self-employed;<br>3: Retired;<br>4: Looking after the home;<br>5: Carer for close family member(s);<br>6: Providing childcare for family;<br>7: Unable to work due to sickness;<br>8: Unemployed;<br>9: Unpaid or voluntary work;<br>10: Student;<br>-1: Do not know;<br>-3: Prefer not to answer | 1–10 |

*Note*. T/Screen = Touchscreen Questionnaire; MHQ = Mental Health Questionnaire; MWB = Mental Well-being Questionnaire. a. Instances where *None of the above* is recorded as 0 to represent the absence of symptoms or activities. b. Special cases where *All of the above* is recoded to match the actual number of available options.

**Table S5. Mental Health Questions Assessed across Questionnaires**

| Mental Health Category | T/Screen 2009+ | | | | MHQ 2016+ | MWB 2022+ | T/Screen 2009+ Questions | Online Questions | Notes on Differences |
|---|---|---|---|---|---|---|---|---|---|
| | 2006–10 | 2012–13 | 2014+ | 2019+ | | | | | |
| Depression | 2060 | 2060 | 2060 | 2060 | 20514 | 29002 | Over the past two weeks, how often have you had little interest or pleasure in doing things? | Over the last 2 weeks, how often have you been bothered *by any of the following problems*… Little interest or pleasure in doing things | Reformatted |
| Depression | 2050 | 2050 | 2050 | 2050 | 20510 | 29003 | Over the past two weeks, how often have you felt down, depressed or hopeless? | … Feeling down, depressed, or hopeless | Reformatted |
| Depression | 2080 | 2080 | 2080 | 2080 | 20519 | 29005 | Over the past two weeks, how often have you felt tired or had little energy? | … Feeling tired or having little energy | Reformatted |
| Depression | 2070 | 2070 | 2070 | 2070 | 20518 | 29009 | Over the past two weeks, how often have you felt tense, fidgety or restless? | … *Moving or speaking so slowly that other people could have noticed?* Or the opposite - being so fidgety or restless that you have been moving around a lot more than usual | Symptoms added |
| Depression | 4598 | 4598 | 4598 | 4598 | 20446 | 29011 | Have you ever had a time when you were feeling depressed or down for *at least a whole week*? | Have you ever had a time in your life when you felt sad, blue, or depressed *for two weeks or more* in a row? | Symptom duration extended |
| Depression | 4631 | 4631 | 4631 | 4631 | 20441 | 29012 | Have you ever had a time when you were uninterested in things or unable to enjoy the things you used to for *at least a whole week*? | Have you ever had a time in your life *lasting two weeks or more* when you lost interest in most things like hobbies, work, or activities that usually give you pleasure? | Symptom duration extended |
| Bipolar Disorder | 4642 | 4642 | 4642 | 4642 | 20501 | 29049 | Have you ever had a period of time *lasting at least two days* when you were feeling so good, “high”, excited or | Have you ever had a period of time when you were feeling so good, “high”, excited or “hyper” that other people thought you were not your | Symptom duration removed |

| Mental Health Category | T/Screen 2009+ | | | | MHQ 2016+ | MWB 2022+ | T/Screen 2009+ Questions | Online Questions | Notes on Differences |
|---|---|---|---|---|---|---|---|---|---|
| | 2006–10 | 2012–13 | 2014+ | 2019+ | | | | | |
| | | | | | | | "hyper" that other people thought you were not your normal self or you were so "hyper" that you got into trouble? | normal self or you were so "hyper" that you got into trouble? | |
| Bipolar Disorder | 4653 | 4653 | 4653 | 4653 | 20502 | 29050 | Have you ever had a period of time *lasting at least two days* when you were so irritable that you found yourself shouting at people or starting fights or arguments? | Have you ever had a period of time when you were so irritable that you found yourself shouting at people or starting fights or arguments? | Symptom duration removed |
| Bipolar Disorder | 6156 | 6156 | 6156 | 6156 | 20548 | 29051 | Please try to remember a period when you were in a "high" or "irritable" state and select which of the following apply. | Please try to remember a period when you were in a "high" or "irritable" state and select which of the following apply. | More symptom options added |
| Bipolar Disorder | 5663 | 5663 | 5663 | 5663 | 20492 | 29052 | What is the longest time period that these "high" or "irritable" periods have lasted? | What is the longest time period that these "high" or "irritable" periods have lasted? | Identical |
| Bipolar Disorder | 5674 | 5674 | 5674 | 5674 | — | 29056 | How much of a problem have these "high" or "irritable" periods caused you? | How much of a problem have these "high" or "irritable" periods caused you… Needed treatment | Reformatted |
| Bipolar Disorder | 5674 | 5674 | 5674 | 5674 | — | 29057 | How much of a problem have these "high" or "irritable" periods caused you? | … Caused problems with work, relationships, finances, the law or other aspects of life | Reformatted |

*Note.* T/Screen = Touchscreen Questionnaire; MHQ = Mental Health Questionnaire; MWB = Mental Well-being Questionnaire.

**Table S6. Cohort Characteristics**

| Variable | Overall (*N* = 8,415) | Discovery (*N* = 4,207) | Validation (*N* = 4,208) | Test Statistics | *p* |
|---|---|---|---|---|---|
| **Demographics** | | | | | |
| Age at initial visit (years) | 62.84 ± 7.36 | 62.90 ± 7.42 | 62.77 ± 7.29 | $t = 0.82$ | .414 |
| Sex, female | 4,515 (53.7%) | 2,262 (53.8%) | 2,253 (53.5%) | $\chi^2 = 0.04$ | .835 |
| **Initial imaging visit (2014–2022)** | | | | | |
| Site, Cheadle | 3,620 (43.0%) | 1,826 (43.4%) | 1,794 (42.6%) | | |
| Site, Newcastle | 4,457 (53.0%) | 2,208 (52.5%) | 2,249 (53.4%) | | |
| Site, Reading | 338 (4.0%) | 173 (4.1%) | 165 (3.9%) | $\chi^2 = 0.85$ | .654 |
| Head size | 1.30 ± 0.12 | 1.30 ± 0.12 | 1.30 ± 0.12 | $t = 1.30$ | .195 |
| Head motion, rfMRI | 0.12 ± 0.05 | 0.12 ± 0.05 | 0.12 ± 0.05 | $t = -0.74$ | .461 |
| Head motion, tfMRI | 0.14 ± 0.06 | 0.14 ± 0.06 | 0.14 ± 0.06 | $t = 0.25$ | .801 |
| Scan date | 2014–2022 | Median day 1,294 [1,040–1,653] | Median day 1,314 [1,055–1,680] | $U = -1.90$ | .057 |
| **Repeat imaging visit (2019–2024)** | | | | | |
| Age at repeat visit (years) | 67.44 ± 7.55 | 67.52 ± 7.59 | 67.36 ± 7.52 | $t = 0.94$ | .346 |
| Site, Cheadle | 3,287 (42.0%) | 1,643 (42.0%) | 1,644 (41.9%) | | |
| Site, Newcastle | 4,206 (53.7%) | 2,093 (53.5%) | 2,113 (53.9%) | | |
| Site, Reading | 338 (4.3%) | 173 (4.4%) | 165 (4.2%) | $\chi^2 = 0.26$ | .877 |
| Head size† | 1.30 ± 0.12 | 1.30 ± 0.12 | 1.30 ± 0.12 | $t = 1.62$ | .105 |
| Head motion, rfMRI | 0.12 ± 0.05 | 0.12 ± 0.06 | 0.12 ± 0.05 | $t = 0.48$ | .634 |
| Head motion, tfMRI | 0.14 ± 0.06 | 0.15 ± 0.06 | 0.14 ± 0.06 | $t = 1.33$ | .185 |
| Scan date | 2019–2024 | Median day 1,534 [284–1,838] | Median day 1,528 [283–1,832] | $U = 0.83$ | .408 |
| Interval between scans (years) | 4.99 [2.31–5.74] | 5.00 [2.32–5.84] | 4.99 [2.31–5.66] | $U = 1.52$ | .128 |
| **Online questionnaires** | | | | | |
| MHQ 2016+ completion date | 2016–2017 | Median day 51 [40–65] | Median day 52 [41–65] | $U = -1.04$ | .299 |
| MWB 2022+ completion date | 2022–2023 | Median day 31 [24–40] | Median day 32 [27–40] | $U = -1.75$ | .081 |

*Note.* Continuous variables presented as mean ± *SD*. Scan and questionnaire completion dates presented as range (overall, in calendar years) or median [Q25–Q75] (discovery and validation, in days since the earliest date in the cohort). Interval between scans presented as median [Q25–Q75] in years throughout. $\chi^2$ = chi-square statistic; *t* = independent-samples *t*-statistic; *U* = standardised Mann-Whitney *U* statistic.

**Table S7. Effect Size and Replicability of Brain-Mental Health Associations by Mental Health Domain and Category**

| Domain / Category | *n* MHPs | *n* Correlations | Median \|*r*\| | Top 1% \|*r*\| | Max \|*r*\| | *p* < .05 (%) | Replication (%) | *q* < .05 (*n*) | Mean \|Δ*r*\| | $r_{between}$ |
|---|---|---|---|---|---|---|---|---|---|---|
| **Mental disorders** | **196** | **3,445,680** | **0.020** | **0.118** | **0.430** | **6.0** | **7.8** | **65** | **0.036** | **0.042** |
| Anxiety | 44 | 773,520 | 0.021 | 0.122 | 0.289 | 6.0 | 7.0 | 0 | 0.038 | 0.043 |
| Bipolar disorder | 14 | 246,120 | 0.024 | 0.135 | 0.291 | 5.1 | 5.0 | 0 | 0.045 | 0.020 |
| Depression | 86 | 1,511,880 | 0.019 | 0.093 | 0.248 | 5.8 | 6.7 | 2 | 0.033 | 0.047 |
| Eating disorders | 15 | 263,700 | 0.033 | 0.177 | 0.430 | 5.4 | 6.0 | 0 | 0.062 | 0.012 |
| Non-specific diagnoses | 9 | 158,220 | 0.015 | 0.076 | 0.219 | 6.8 | 9.4 | 0 | 0.025 | 0.128 |
| PTSD | 5 | 87,900 | 0.016 | 0.083 | 0.169 | 5.8 | 6.1 | 0 | 0.028 | 0.089 |
| Substance use | 23 | 404,340 | 0.018 | 0.093 | 0.232 | 6.9 | 14.5 | 244 | 0.031 | 0.081 |
| **Transdiagnostic features** | **60** | **1,054,800** | **0.015** | **0.074** | **0.325** | **7.0** | **12.3** | **27** | **0.024** | **0.141** |
| General health | 5 | 87,900 | 0.015 | 0.062 | 0.123 | 7.0 | 9.7 | 1 | 0.023 | 0.181 |
| Neuroticism | 26 | 457,080 | 0.015 | 0.064 | 0.135 | 6.9 | 13.0 | 42 | 0.023 | 0.166 |
| Resilience | 6 | 105,480 | 0.014 | 0.061 | 0.124 | 6.8 | 13.6 | 0 | 0.023 | 0.172 |
| Self-harm behaviours | 6 | 105,480 | 0.019 | 0.130 | 0.325 | 6.7 | 6.3 | 0 | 0.036 | 0.020 |
| Subjective well-being | 17 | 298,860 | 0.015 | 0.064 | 0.161 | 7.5 | 13.5 | 2 | 0.023 | 0.190 |
| **Environmental factors** | **53** | **931,740** | **0.015** | **0.072** | **0.227** | **6.7** | **8.9** | **18** | **0.025** | **0.079** |
| Adverse experiences | 30 | 527,400 | 0.014 | 0.062 | 0.160 | 6.4 | 7.0 | 0 | 0.024 | 0.069 |
| COVID exposure | 4 | 70,320 | 0.019 | 0.111 | 0.224 | 6.0 | 9.8 | 0 | 0.035 | 0.011 |
| Social factors | 19 | 334,020 | 0.015 | 0.075 | 0.227 | 7.4 | 11.5 | 99 | 0.025 | 0.118 |

*Note.* IDP = imaging-derived phenotype; MHP = mental health phenotype. Categories are those retained after phenotype selection; behavioural addiction and psychotic experiences yielded no phenotypes. Effect-size and significance statistics are from the discovery subset ($N$ = 4,207); validation-subset values agreed to three decimal places throughout and are omitted. Median $|r|$, top 1% $|r|$ and max $|r|$ summarise the distribution of absolute Pearson correlations. $p < .05$ (%) is the percentage of associations significant before correction; $q < .05$ ($n$) is the number surviving FDR correction applied within each row. Replication (%) is the percentage of discovery associations with $p < .05$ that were also significant at $p < .05$ in the validation subset ($N = 4{,}208$). Mean $|\Delta r|$ is the mean absolute difference in effect size between subsets, and $r_{between}$ is the correlation between discovery and validation effect sizes across all associations in that row.

**Table S8. Effect Size and Replicability of Brain-Mental Health Associations by Imaging Modality**

| Modality | *n* IDPs | *n* Correlations | Median \|*r*\| | Top 1% \|*r*\| | Max \|*r*\| | *p* < .05 (%) | Replication (%) | *q* < .05 (*n*) | Mean \|Δ*r*\| | $r_{between}$ |
|---|---|---|---|---|---|---|---|---|---|---|
| ASL | 50 | 15,450 | 0.038 | 0.232 | 0.430 | 6.4 | 9.7 | 0 | 0.066 | 0.121 |
| SWI | 64 | 19,776 | 0.019 | 0.109 | 0.223 | 7.1 | 18.4 | 0 | 0.033 | 0.151 |
| T1 | 2,864 | 884,976 | 0.017 | 0.108 | 0.316 | 6.0 | 9.1 | 163 | 0.032 | 0.072 |
| T2 | 6 | 1,854 | 0.021 | 0.120 | 0.221 | 11.9 | 14.6 | 2 | 0.030 | 0.287 |
| dMRI | 1,350 | 417,150 | 0.018 | 0.109 | 0.288 | 7.2 | 9.4 | 0 | 0.032 | 0.130 |
| rfMRI | 13,182 | 4,073,238 | 0.018 | 0.107 | 0.350 | 6.3 | 8.8 | 24 | 0.032 | 0.043 |
| tfMRI | 64 | 19,776 | 0.020 | 0.123 | 0.238 | 8.5 | 9.3 | 0 | 0.035 | 0.024 |

*Note.* Each modality was correlated with all 309 MHPs. ASL IDPs were available at the repeat imaging visit only. Column definitions are as in Table S7.

**Table S9. Effect Size and Replicability of Brain-Mental Health Associations by Mental Health Category and Imaging Modality**

| Category / Modality | $n$ Correlations | Median $\lvert r\rvert$ | Top 1% $\lvert r\rvert$ | Max $\lvert r\rvert$ | $p < .05$ (%) | Replication (%) | $q < .05$ ($n$) | Mean $\lvert\Delta r\rvert$ | $r_{between}$ |
|---|---|---|---|---|---|---|---|---|---|
| **Anxiety** | | | | | | | | | |
| ASL | 2,200 | 0.042 | 0.177 | 0.289 | 4.5 | 7.0 | 0 | 0.086 | -0.136 |
| SWI | 2,816 | 0.021 | 0.120 | 0.183 | 4.1 | 6.0 | 0 | 0.039 | 0.030 |
| T1 | 126,016 | 0.020 | 0.127 | 0.259 | 6.1 | 5.5 | 0 | 0.038 | 0.046 |
| T2 | 264 | 0.027 | 0.204 | 0.221 | 15.9 | 4.8 | 0 | 0.033 | 0.381 |
| dMRI | 59,400 | 0.022 | 0.121 | 0.232 | 7.8 | 7.0 | 0 | 0.037 | 0.184 |
| rfMRI | 580,008 | 0.021 | 0.120 | 0.277 | 5.7 | 7.3 | 0 | 0.038 | 0.027 |
| tfMRI | 2,816 | 0.024 | 0.139 | 0.195 | 8.2 | 9.9 | 0 | 0.042 | 0.028 |
| **Bipolar disorder** | | | | | | | | | |
| ASL | 700 | 0.042 | 0.204 | 0.258 | 1.9 | 0.0 | 0 | 0.096 | -0.286 |
| SWI | 896 | 0.028 | 0.151 | 0.212 | 6.7 | 23.3 | 0 | 0.046 | 0.229 |
| T1 | 40,096 | 0.024 | 0.130 | 0.291 | 4.8 | 4.7 | 0 | 0.045 | 0.025 |
| T2 | 84 | 0.022 | 0.128 | 0.128 | 6.0 | 0.0 | 0 | 0.041 | 0.120 |
| dMRI | 18,900 | 0.024 | 0.126 | 0.240 | 5.6 | 4.1 | 0 | 0.047 | 0.009 |
| rfMRI | 184,548 | 0.024 | 0.136 | 0.286 | 5.1 | 5.0 | 0 | 0.044 | 0.025 |
| tfMRI | 896 | 0.029 | 0.186 | 0.238 | 9.0 | 6.2 | 0 | 0.057 | -0.172 |
| **Depression** | | | | | | | | | |
| ASL | 4,300 | 0.037 | 0.183 | 0.248 | 5.7 | 20.3 | 0 | 0.065 | 0.080 |
| SWI | 5,504 | 0.021 | 0.094 | 0.167 | 7.4 | 16.4 | 0 | 0.033 | 0.151 |

| Category / Modality | $n$ Correlations | Median $\|r\|$ | Top 1% $\|r\|$ | Max $\|r\|$ | $p < .05$ (%) | Replication (%) | $q < .05$ ($n$) | Mean $\|\Delta r\|$ | $r_{between}$ |
|---|---|---|---|---|---|---|---|---|---|
| T1 | 246,304 | 0.018 | 0.090 | 0.214 | 5.4 | 7.1 | 0 | 0.032 | 0.058 |
| T2 | 516 | 0.019 | 0.081 | 0.097 | 9.1 | 10.6 | 0 | 0.027 | 0.280 |
| dMRI | 116,100 | 0.019 | 0.092 | 0.211 | 6.2 | 6.8 | 0 | 0.031 | 0.137 |
| rfMRI | 1,133,652 | 0.019 | 0.092 | 0.222 | 5.8 | 6.5 | 3 | 0.033 | 0.035 |
| tfMRI | 5,504 | 0.021 | 0.125 | 0.171 | 9.2 | 8.7 | 0 | 0.037 | -0.015 |
| **Eating disorders** | | | | | | | | | |
| ASL | 750 | 0.086 | 0.393 | 0.430 | 18.1 | 4.4 | 0 | 0.104 | 0.542 |
| SWI | 960 | 0.036 | 0.162 | 0.223 | 5.2 | 6.0 | 0 | 0.060 | 0.067 |
| T1 | 42,960 | 0.032 | 0.172 | 0.316 | 4.8 | 4.7 | 0 | 0.058 | 0.091 |
| T2 | 90 | 0.037 | 0.165 | 0.167 | 7.8 | 0.0 | 0 | 0.056 | 0.081 |
| dMRI | 20,250 | 0.037 | 0.179 | 0.288 | 9.5 | 7.1 | 2 | 0.065 | 0.101 |
| rfMRI | 197,730 | 0.033 | 0.176 | 0.350 | 5.1 | 6.1 | 0 | 0.062 | -0.031 |
| tfMRI | 960 | 0.030 | 0.141 | 0.178 | 3.0 | 0.0 | 0 | 0.061 | 0.030 |
| **Non-specific diagnoses** | | | | | | | | | |
| ASL | 450 | 0.060 | 0.153 | 0.219 | 22.9 | 6.8 | 12 | 0.068 | 0.006 |
| SWI | 576 | 0.020 | 0.085 | 0.131 | 15.8 | 13.2 | 7 | 0.027 | 0.172 |
| T1 | 25,776 | 0.014 | 0.070 | 0.162 | 4.9 | 3.9 | 0 | 0.024 | -0.013 |
| T2 | 54 | 0.013 | 0.061 | 0.062 | 0.0 | 0.0 | 0 | 0.024 | 0.346 |
| dMRI | 12,150 | 0.017 | 0.086 | 0.153 | 8.1 | 10.7 | 0 | 0.024 | 0.267 |
| rfMRI | 118,638 | 0.015 | 0.074 | 0.162 | 6.9 | 10.0 | 0 | 0.025 | 0.137 |
| tfMRI | 576 | 0.026 | 0.104 | 0.129 | 22.7 | 14.5 | 0 | 0.037 | 0.347 |

| Category / Modality | *n* Correlations | Median \|*r*\| | Top 1% \|*r*\| | Max \|*r*\| | *p* < .05 (%) | Replication (%) | *q* < .05 (*n*) | Mean \|Δ*r*\| | $r_{\text{between}}$ |
|---|---|---|---|---|---|---|---|---|---|
| **PTSD** | | | | | | | | | |
| ASL | 250 | 0.033 | 0.127 | 0.146 | 2.8 | 0.0 | 0 | 0.060 | 0.042 |
| SWI | 320 | 0.018 | 0.090 | 0.125 | 9.4 | 13.3 | 0 | 0.026 | 0.379 |
| T1 | 14,320 | 0.016 | 0.083 | 0.146 | 5.6 | 6.4 | 0 | 0.026 | 0.104 |
| T2 | 30 | 0.036 | 0.072 | 0.072 | 20.0 | 50.0 | 0 | 0.016 | 0.846 |
| dMRI | 6,750 | 0.014 | 0.071 | 0.121 | 3.3 | 7.3 | 0 | 0.029 | -0.003 |
| rfMRI | 65,910 | 0.017 | 0.083 | 0.169 | 6.1 | 5.9 | 0 | 0.028 | 0.092 |
| tfMRI | 320 | 0.019 | 0.080 | 0.096 | 6.9 | 0.0 | 0 | 0.028 | 0.015 |
| **Substance use** | | | | | | | | | |
| ASL | 1,150 | 0.040 | 0.139 | 0.232 | 7.0 | 21.3 | 0 | 0.049 | 0.327 |
| SWI | 1,472 | 0.024 | 0.124 | 0.200 | 15.6 | 39.3 | 41 | 0.033 | 0.424 |
| T1 | 65,872 | 0.019 | 0.094 | 0.214 | 9.3 | 21.9 | 529 | 0.031 | 0.061 |
| T2 | 138 | 0.016 | 0.081 | 0.085 | 5.8 | 25.0 | 0 | 0.034 | 0.185 |
| dMRI | 31,050 | 0.018 | 0.093 | 0.174 | 8.4 | 27.0 | 20 | 0.031 | 0.136 |
| rfMRI | 303,186 | 0.018 | 0.091 | 0.229 | 6.2 | 10.0 | 0 | 0.031 | 0.057 |
| tfMRI | 1,472 | 0.018 | 0.088 | 0.127 | 7.7 | 14.9 | 0 | 0.030 | 0.256 |
| **General health** | | | | | | | | | |
| ASL | 250 | 0.036 | 0.104 | 0.123 | 3.6 | 0.0 | 0 | 0.049 | -0.200 |
| SWI | 320 | 0.020 | 0.061 | 0.070 | 10.9 | 54.3 | 2 | 0.023 | 0.431 |
| T1 | 14,320 | 0.016 | 0.064 | 0.108 | 9.0 | 10.4 | 1 | 0.023 | 0.218 |
| T2 | 30 | 0.026 | 0.062 | 0.062 | 26.7 | 37.5 | 0 | 0.028 | 0.395 |

| Category / Modality | *n* Correlations | Median \|*r*\| | Top 1% \|*r*\| | Max \|*r*\| | *p* < .05 (%) | Replication (%) | *q* < .05 (*n*) | Mean \|Δ*r*\| | $r_{between}$ |
|---|---|---|---|---|---|---|---|---|---|
| dMRI | 6,750 | 0.015 | 0.064 | 0.107 | 9.1 | 16.9 | 1 | 0.022 | 0.329 |
| rfMRI | 65,910 | 0.014 | 0.061 | 0.108 | 6.4 | 8.0 | 0 | 0.022 | 0.144 |
| tfMRI | 320 | 0.015 | 0.059 | 0.068 | 6.9 | 18.2 | 0 | 0.029 | -0.241 |
| **Neuroticism** | | | | | | | | | |
| ASL | 1,300 | 0.025 | 0.093 | 0.126 | 2.0 | 7.7 | 0 | 0.048 | -0.188 |
| SWI | 1,664 | 0.015 | 0.056 | 0.070 | 5.5 | 6.5 | 0 | 0.024 | 0.105 |
| T1 | 74,464 | 0.014 | 0.060 | 0.102 | 6.1 | 9.7 | 0 | 0.023 | 0.132 |
| T2 | 156 | 0.016 | 0.069 | 0.072 | 10.9 | 0.0 | 0 | 0.023 | -0.012 |
| dMRI | 35,100 | 0.014 | 0.060 | 0.117 | 5.2 | 5.7 | 0 | 0.022 | 0.127 |
| rfMRI | 342,732 | 0.015 | 0.065 | 0.135 | 7.3 | 14.2 | 54 | 0.023 | 0.179 |
| tfMRI | 1,664 | 0.015 | 0.062 | 0.082 | 4.6 | 9.2 | 0 | 0.023 | -0.001 |
| **Resilience** | | | | | | | | | |
| ASL | 300 | 0.030 | 0.088 | 0.112 | 2.0 | 33.3 | 0 | 0.050 | -0.307 |
| SWI | 384 | 0.012 | 0.062 | 0.075 | 3.1 | 0.0 | 0 | 0.021 | -0.131 |
| T1 | 17,184 | 0.013 | 0.057 | 0.095 | 5.0 | 5.7 | 0 | 0.021 | 0.136 |
| T2 | 36 | 0.017 | 0.064 | 0.064 | 11.1 | 0.0 | 0 | 0.035 | -0.662 |
| dMRI | 8,100 | 0.014 | 0.064 | 0.103 | 6.3 | 8.3 | 0 | 0.022 | 0.302 |
| rfMRI | 79,092 | 0.015 | 0.061 | 0.124 | 7.2 | 15.3 | 0 | 0.023 | 0.171 |
| tfMRI | 384 | 0.023 | 0.062 | 0.072 | 13.0 | 4.0 | 0 | 0.026 | 0.107 |
| **Self-harm behaviours** | | | | | | | | | |
| ASL | 300 | 0.070 | 0.280 | 0.325 | 17.7 | 0.0 | 0 | 0.110 | -0.562 |

| Category / Modality | *n* Correlations | Median $|r|$ | Top 1% $|r|$ | Max $|r|$ | $p < .05$ (%) | Replication (%) | $q < .05$ ($n$) | Mean $|\Delta r|$ | $r_{between}$ |
|---|---|---|---|---|---|---|---|---|---|
| SWI | 384 | 0.018 | 0.095 | 0.111 | 3.4 | 15.4 | 0 | 0.035 | 0.050 |
| T1 | 17,184 | 0.018 | 0.128 | 0.239 | 5.3 | 11.2 | 0 | 0.035 | 0.046 |
| T2 | 36 | 0.011 | 0.083 | 0.083 | 0.0 | 0.0 | 0 | 0.026 | 0.225 |
| dMRI | 8,100 | 0.018 | 0.120 | 0.210 | 6.4 | 6.6 | 0 | 0.036 | -0.045 |
| rfMRI | 79,092 | 0.019 | 0.130 | 0.300 | 7.1 | 5.5 | 0 | 0.036 | 0.030 |
| tfMRI | 384 | 0.021 | 0.112 | 0.143 | 6.5 | 4.0 | 0 | 0.039 | -0.166 |
| **Subjective well-being** | | | | | | | | | |
| ASL | 850 | 0.035 | 0.109 | 0.161 | 5.8 | 0.0 | 0 | 0.057 | -0.234 |
| SWI | 1,088 | 0.016 | 0.078 | 0.102 | 8.2 | 22.5 | 3 | 0.025 | 0.247 |
| T1 | 48,688 | 0.015 | 0.064 | 0.123 | 7.3 | 14.8 | 2 | 0.023 | 0.187 |
| T2 | 102 | 0.029 | 0.104 | 0.105 | 29.4 | 33.3 | 16 | 0.035 | 0.132 |
| dMRI | 22,950 | 0.015 | 0.067 | 0.109 | 7.3 | 8.6 | 0 | 0.022 | 0.158 |
| rfMRI | 224,094 | 0.015 | 0.063 | 0.121 | 7.5 | 13.6 | 1 | 0.023 | 0.195 |
| tfMRI | 1,088 | 0.015 | 0.053 | 0.061 | 4.0 | 25.0 | 0 | 0.019 | 0.276 |
| **Adverse experiences** | | | | | | | | | |
| ASL | 1,500 | 0.035 | 0.123 | 0.160 | 6.1 | 4.4 | 0 | 0.050 | 0.182 |
| SWI | 1,920 | 0.014 | 0.061 | 0.092 | 5.1 | 9.2 | 0 | 0.025 | 0.063 |
| T1 | 85,920 | 0.014 | 0.060 | 0.107 | 5.8 | 7.6 | 0 | 0.024 | 0.054 |
| T2 | 180 | 0.019 | 0.055 | 0.059 | 8.9 | 43.8 | 0 | 0.022 | 0.361 |
| dMRI | 40,500 | 0.016 | 0.062 | 0.113 | 8.9 | 9.7 | 0 | 0.025 | 0.146 |
| rfMRI | 395,460 | 0.014 | 0.062 | 0.118 | 6.2 | 6.5 | 0 | 0.023 | 0.061 |

| Category / Modality | *n* Correlations | Median \|*r*\| | Top 1% \|*r*\| | Max \|*r*\| | *p* < .05 (%) | Replication (%) | *q* < .05 (*n*) | Mean \|Δ*r*\| | $r_{between}$ |
|---|---|---|---|---|---|---|---|---|---|
| tfMRI | 1,920 | 0.018 | 0.066 | 0.082 | 11.8 | 5.7 | 9 | 0.025 | 0.110 |
| **COVID exposure** | | | | | | | | | |
| ASL | 200 | 0.039 | 0.132 | 0.142 | 4.0 | 0.0 | 0 | 0.056 | -0.030 |
| SWI | 256 | 0.020 | 0.115 | 0.121 | 7.8 | 5.0 | 0 | 0.031 | 0.101 |
| T1 | 11,456 | 0.020 | 0.121 | 0.187 | 7.1 | 2.3 | 0 | 0.034 | 0.031 |
| T2 | 24 | 0.017 | 0.156 | 0.156 | 25.0 | 0.0 | 2 | 0.041 | 0.414 |
| dMRI | 5,400 | 0.018 | 0.116 | 0.197 | 4.8 | 13.0 | 0 | 0.038 | -0.149 |
| rfMRI | 52,728 | 0.019 | 0.107 | 0.224 | 5.9 | 11.6 | 0 | 0.035 | 0.025 |
| tfMRI | 256 | 0.022 | 0.082 | 0.088 | 9.0 | 0.0 | 0 | 0.036 | 0.107 |
| **Social factors** | | | | | | | | | |
| ASL | 950 | 0.033 | 0.128 | 0.177 | 5.9 | 0.0 | 0 | 0.049 | 0.035 |
| SWI | 1,216 | 0.015 | 0.064 | 0.136 | 4.8 | 6.9 | 0 | 0.024 | 0.155 |
| T1 | 54,416 | 0.014 | 0.070 | 0.171 | 5.4 | 6.8 | 0 | 0.025 | 0.045 |
| T2 | 114 | 0.022 | 0.105 | 0.132 | 21.1 | 0.0 | 3 | 0.028 | 0.245 |
| dMRI | 25,650 | 0.016 | 0.082 | 0.174 | 9.5 | 7.7 | 0 | 0.025 | 0.197 |
| rfMRI | 250,458 | 0.015 | 0.074 | 0.227 | 7.7 | 12.7 | 117 | 0.025 | 0.121 |
| tfMRI | 1,216 | 0.018 | 0.075 | 0.135 | 7.3 | 11.2 | 0 | 0.023 | 0.273 |

*Note.* Categories are ordered by mental health domain (mental disorders, transdiagnostic features, environmental factors). Column definitions are as in Table S7.

**Table S10. Effect Size and Replicability of Brain-Mental Health Associations Across Temporal Configurations**

| Configuration | *n* IDPs | *n* MHPs | *n* Correlations | Median $\|r\|$ | Top 1% $\|r\|$ | Max $\|r\|$ | $p < .05$ (%) | Replication (%) | $q < .05$ (*n*) | Mean $\|\Delta r\|$ | $r_{\text{between}}$ |
|---|---|---|---|---|---|---|---|---|---|---|---|
| **Same-day** | | | | | | | | | | | |
| IDP V1 × MHP V1 | 8,765 | 36 | 315,540 | 0.013 | 0.055 | 0.123 | 7.5 | 13.6 | 73 | 0.020 | 0.175 |
| IDP V2 × MHP V2 | 8,765 | 36 | 315,540 | 0.019 | 0.086 | 0.218 | 6.4 | 9.2 | 1 | 0.031 | 0.108 |
| **Cross-visit** | | | | | | | | | | | |
| IDP V1 × MHP V2 | 8,765 | 36 | 315,540 | 0.013 | 0.057 | 0.162 | 7.3 | 13.1 | 69 | 0.020 | 0.161 |
| IDP V2 × MHP V1 | 8,765 | 36 | 315,540 | 0.019 | 0.081 | 0.185 | 6.0 | 7.6 | 3 | 0.031 | 0.094 |

*Note.* Analyses are restricted to participants attending both imaging visits and to the 8,765 IDPs and 36 MHPs available at both, so ASL IDPs are excluded. Same-day configurations pair brain and mental health measures from the same visit; cross-visit configurations pair them across visits. Column definitions are as in Table S7.

**Table S11. Effect Size and Replicability of Within-Subject Repeated-Measures Associations**

| MHP set | *n* IDPs | *n* MHPs | *n* Correlations | Median $\|r_{rm}\|$ | Top 1% $\|r_{rm}\|$ | Max $\|r_{rm}\|$ | $p < .05$ (%) | Replication (%) | $q < .05$ (*n*) | Mean $\|\Delta r_{rm}\|$ | $r_{\text{between}}$ |
|---|---|---|---|---|---|---|---|---|---|---|---|
| **All MHPs** | **8,765** | **36** | **315,540** | **0.019** | **0.104** | **0.252** | **5.0** | **5.2** | **0** | **0.034** | **−0.004** |
| Trait-like | 8,765 | 22 | 192,830 | 0.020 | 0.118 | 0.252 | 5.3 | 5.0 | 0 | 0.037 | −0.010 |
| State-like | 8,765 | 14 | 122,710 | 0.018 | 0.066 | 0.118 | 4.6 | 5.6 | 0 | 0.029 | 0.015 |

*Note.* $r_{rm}$ = repeated-measures correlation, computed within participants across the two imaging visits. Trait-like MHPs are lifetime measures and state-like MHPs are current-state measures; the two sets are mutually exclusive and together comprise all 36 MHPs. Restrictions and column definitions are as in Table S10.
